# AdventML: Advanced Enzyme Temperature Prediction with Transformer-Based Embeddings and Resampling Strategies

**DOI:** 10.64898/2026.05.30.728975

**Authors:** Jaldert François, Bart De Moor, Vera van Noort

## Abstract

Accurate prediction of enzymes’ optimal catalytic temperature (*T*_opt_) is crucial in biotechnology, as enzymes with extreme *T*_opt_ values are highly desirable for reactions at extreme temperatures and for their general stability. However, experimental determination of *T*_opt_ is costly, labor-intensive, and time-consuming. Meanwhile, existing computational methods suffer from small and imbalanced datasets, suboptimal predictions at extreme temperatures, and insufficient validation.

In this study, we address these challenges by expanding the *T*_opt_ dataset and validating on an independent test set based on sequence similarity. We further tackle these limitations by comparing multiple resampling techniques to improve predictions at extremes and by considering diverse protein representations and multiple machine learning architectures. Overall, the best performing models reached *R*^2^ ≈ 0.64 with MAE ≈ 7–8 °C, while extreme resampling improved tail performance (reducing tail MAE by up to ~1.8 °C). Notably, our models show improved performance over state-of-the-art prediction models. We also demonstrate that accurate prediction of *T*_opt_ is achievable even in the absence of organism growth temperature (OGT). Our *T*_opt_ prediction models are made freely available as AdventML on GitHub.

## 1. Introduction

Enzymes that are stable and active at high temperatures are widely used in pharmaceuticals, food processing, and chemical manufacturing^1–8^, and widely used in molecular biology and protein engineering^9^. However, experimental determination of an enzyme’s optimal catalytic temperature (*T*_opt_) is costly, labor intensive, and time-consuming, highlighting the need for computational prediction methods.

Organism growth temperature (OGT) is often used to estimate protein thermostability and, by extension, the enzyme’s optimal catalytic temperature (*T*_opt_)^9,10^. However, OGT presents significant limitations for predicting enzyme performance at specific temperatures. Firstly, OGT shows only a weak correlation of 0.48 to *T*_opt_ for single enzymes^11^. Secondly, protein thermostability does not always reflect cat-alytic activity at high temperatures^9^. Thirdly, OGT data is not only lacking for many organisms but is also experimentally expensive to determine^9,12^ and is intractable for unculturable organisms. Moreover, many enzymes exhibit *T*_opt_ values substantially higher or lower than their organism’s OGT, making OGT merely a rough estimate. Both the need for better temperature estimates than OGT, and the cost and time related to experimental OGT or *T*_opt_ determination underline the need for accurate enzyme temperature prediction models^13^.

Thermal adaptation is primarily reflected at the protein level^14^ for unicellular organisms (animals have physiological adaptations). Numerous genomic and proteomic features correlate with OGT or *T*_opt_, including dinucleotide pres-ence/absence^15^, GC content in structural RNAs^16^, amino acid composition of the proteome^14,17^, IVYWREL correlation with OGT for Archaea and Bacteria^9,17^, protein length^18^, amino acid properties^17,19,20^, and structural features^21–23^. However, these factors are individually weak determinants of protein unfolding^24,25^ and show even smaller effects on catalytic temperature^9^. Nevertheless, both the link between proteins and temperature, and existing temperature prediction models suggest that building accurate prediction models for *T*_opt_ based on protein characteristics is feasible.

Indeed, recent developments include *T*_opt_ prediction models both in the presence and absence of OGT, showing promising results. Nevertheless, existing approaches remain limited. Models like TOME and TOMER both require OGT data for *T*_opt_ prediction, severely limiting their applications^9,10^. Additionally, a majority of research targets specific enzyme classes, such as Preoptem^26^. Recent improvements in *T*_opt_ prediction include work by Zhang et al. developing a Segment Transformer model, and Seq2Topt by Qiu et al. using ESM2-8M or Progen2-small with fine-tuning^26,27^. However, prediction at temperature extremes remains challenging, and research favors complex deep learning models, often without establishing simple baselines. Critical questions remain regarding model generalizability and out-of-distribution performance, particularly the accuracy at extreme temperatures, that are after all of greatest interest.

In this work, we address these challenges by systematically examining *T*_opt_ prediction across multiple dimensions: protein sequence representations, model complexity, data re-sampling strategies, and training-test similarity. We trained and compared both classical machine learning methods with deep learning based approaches. Surprisingly, we show that a mix of advanced and classical methods is optimal. We demonstrate that accurate *T*_opt_ prediction in the absence of OGT is achievable with recent protein language models (PLMs), while classical physicochemical properties alone prove insufficient. Although resampling methods showed mixed results overall, they provide some improvements at temperature extremes. To facilitate *T*_opt_ prediction for the research community, we release AdventML as both a web portal and an installable GitHub tool.

## 2. Results

The central questions in this research is whether *T*_opt_ prediction is possible in the absence of organism growth temperature (OGT) data and, if so, whether strategies tailored to extreme temperatures can improve the identification of thermostable and cold-adapted enzymes. To this end, we evaluate prediction performance with and without OGT across multiple protein representations, and we extend this comparison to the full training set of ~9, 000 sequences. We further investigate extreme-regression strategies including resampling, bagging and apply loss weighting, to enhance performance in the cold and hot tails of the *T*_opt_ distribution. Finally, we compare several machine learning architectures including deep learning against classical machine learning.

### 2.1 Modest OGT benefit for *T*_opt_ prediction in state-of-the-art PLMs

Many thermostability adapted enzymes have been discovered in, or originate from, temperature resistant organisms. In this context, the optimal growth temperature (OGT) can serve as a useful proxy for assessing enzyme thermostability. However, this relationship is not perfect: not all enzymes from thermophilic hosts are thermostable themselves, and conversely, thermostable enzymes can also arise from mesophilic (and even psychrophilic) organisms. Here, we investigate the extent to which OGT contributes to predictive performance when coupled with modern protein representations derived from protein language models (PLMs) using the original TOMER training-test split. We evaluate both (i) traditional protein variables, such as physicochemical descriptors, and (ii) PLM based embeddings, using Histogram Gradient Boosting, CatBoost, and support vector regression (SVR) with hyperparameter tuning.

Overall, learned protein representations substantially out-perform classical physicochemical descriptors in predictive performance (Fig. 3). Notably, the performance gap between models trained with versus without OGT is small for state-of-the-art PLMs. The difference was smallest for ESMC, ESM3 and PRIME models. This difference, which is near zero for PRIME, was expected since this model is trained to represent and distinguish protein thermostability^28^. Performance for top models, including ESM models and ProtTrans T5, was similar and hovered around *R*^2^ ~ 0.50-0.60 with an *R*^2^ = 0.53 and MAE = 9.44°C for ESM3 SM open and an *R*^2^ = 0.52 with MAE = 9.60°C for ProtT5-XL-50 in the absence of OGT (Supplementary Table S1). Furthermore, the strongest relative improvement occurred for physicochemical features indicating that OGT can partially compensate for the limited sequence derived temperature signal for in these protein representations.

**Figure 1.**
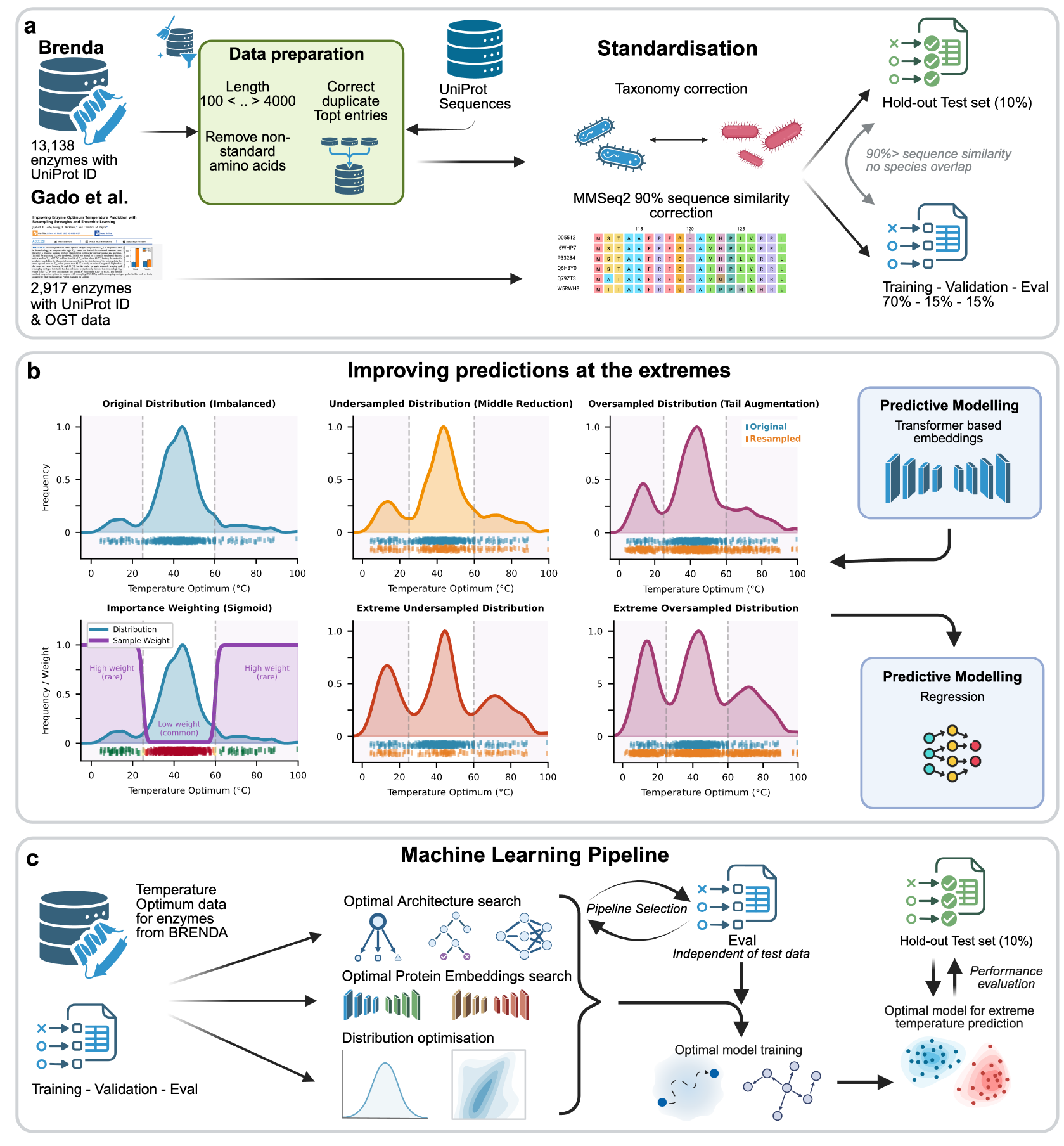
AdventML workflow for *T*_opt_ prediction. (**a**) Data collection and preprocessing from the BRENDA enzyme database, including quality filtering, deduplication, and taxonomic annotation. (**b**) Resampling strategies to improve predictions at temperature extremes. Both oversampling and undersampling approaches were evaluated, including extreme variants that target values *<* 25 ° C and *>* 60 ° C for oversampling, and *>* 25 ° C and *<* 60 °C for undersampling, to balance training examples across the temperature range. (**c**) Systematic search for optimal machine learning strategy, including protein representation comparison, data balancing evaluation, and architecture comparison, with final assessment on an independent held-out test set with sequence and phylogeny similarity correction.

**Figure 2.**
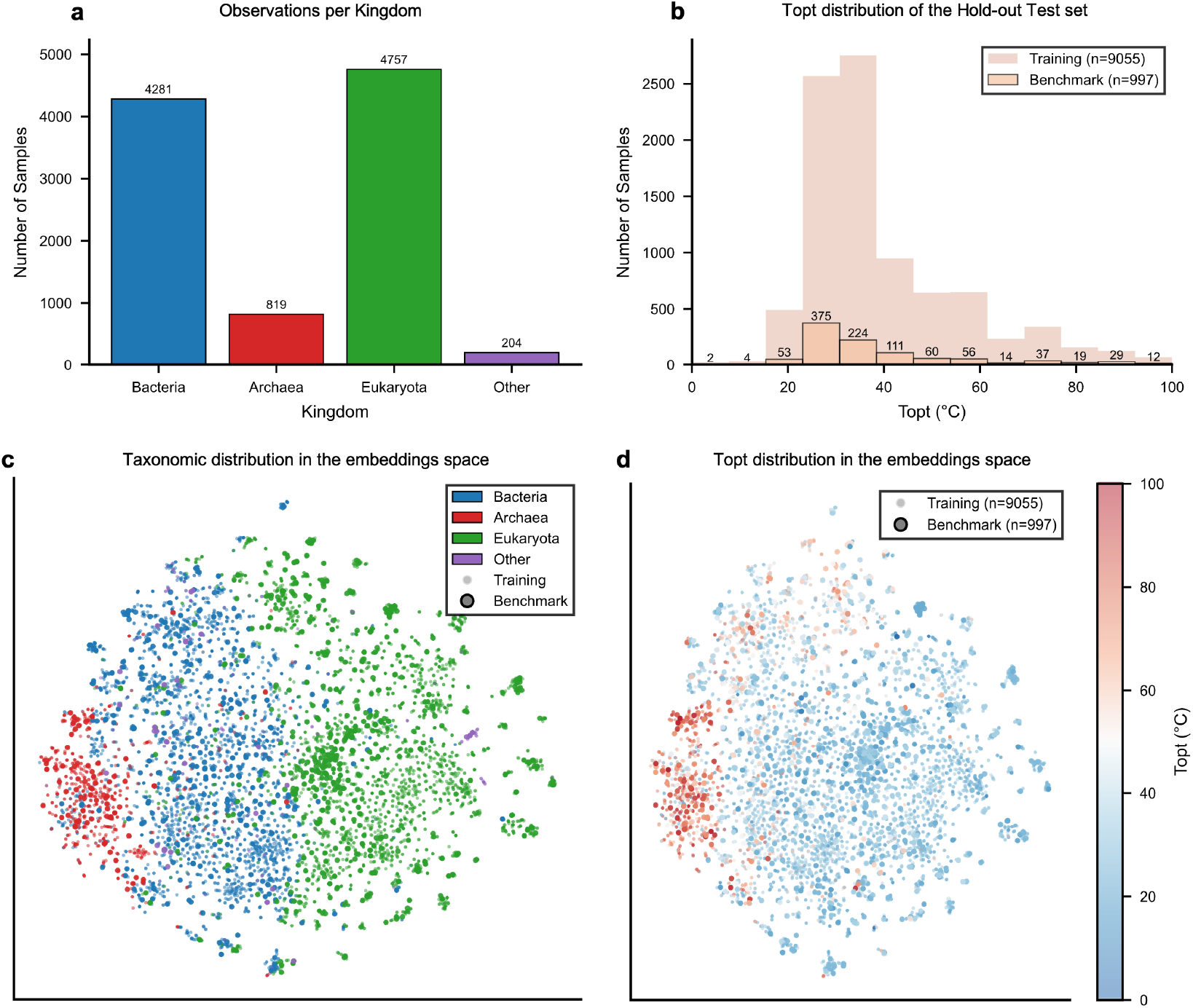
Phylogenetic and thermal diversity of the *T*_opt_ dataset. (**a**) Taxonomic distribution of enzyme sequences across kingdoms in the filtered BRENDA dataset. (**b**) Distribution of optimal temperature (*T*_opt_) values in the training and held-out test sets, showing concentration in the mesophilic range (30–50 ° C) with sparse representation at temperature extremes. (**c**) t-SNE projection of ProtTrans-T5-XL-U50 embeddings colored by taxonomic kingdom, revealing distinct clustering by phylogenetic origin. (**d**) t-SNE projection of ProtTrans-T5-XL-U50 embeddings colored by *T*_opt_, demonstrating a smooth temperature gradient across the representation space that aligns with taxonomic structure.

**Figure 3.**
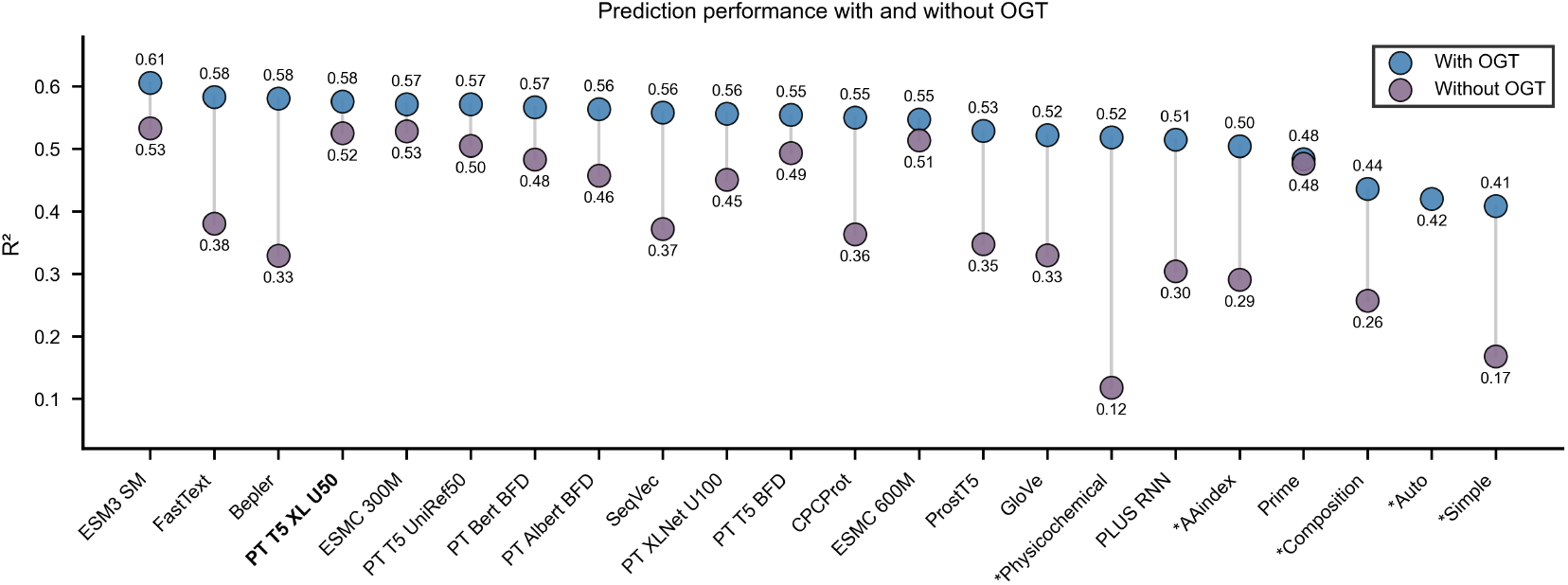
Impact of organism growth temperature on *T*_opt_ prediction across protein representations. Embeddings were computed for the TOMER dataset (*n* = 2,917 sequences). For each representation, Histogram Gradient Boosting, CatBoost, and Support Vector Regression models were trained with hyperparameters optimized via Optuna, and performance was averaged across the three regressors. Modern protein language model embeddings show minimal performance gain from OGT inclusion, whereas classical physicochemical descriptors benefit substantially.

### 2.2 Marginal differences between state-of-the-art protein representations for *T*_opt_ prediction

We benchmarked a broad set of protein representations to identify the most effective feature space for enzyme temperature prediction. In addition to the models described in section 2.1, we expanded the panel of protein language models (PLMs) to include newer transformer based encoders, notably Ankh3 and ESM-2. Furthermore, models were evaluated across a diverse suite of regression models: Histogram Gradient Boosting, XGBoost, CatBoost, QuantileGBR, ElasticNet, support vector regression (SVR), and Kernel Ridge regression. All models were tuned and evaluated on the expanded BRENDA dataset.

Across all regressors, transformer derived embeddings consistently delivered the strongest predictive performance (Fig. 4). These PLM representations substantially outperformed classical physicochemical property descriptors and also improved performance over older learned representations based on convolutional neural networks (CNNs) or recurrent neural networks (RNNs). The best overall results were obtained with ProtT5-XL-U50, which achieved *R*^2^ ≈ 0.53 with MAE = 7.93°C (Supplementary Table S2). While ProtT5-XL-U50 was the top performer, differences among the leading state-of-the-art protein representations were generally small, with most high performing PLMs achieving comparable performance. In contrast, physicochemical property based baselines performed substantially worse, underscoring the importance of current PLMs.

**Figure 4.**
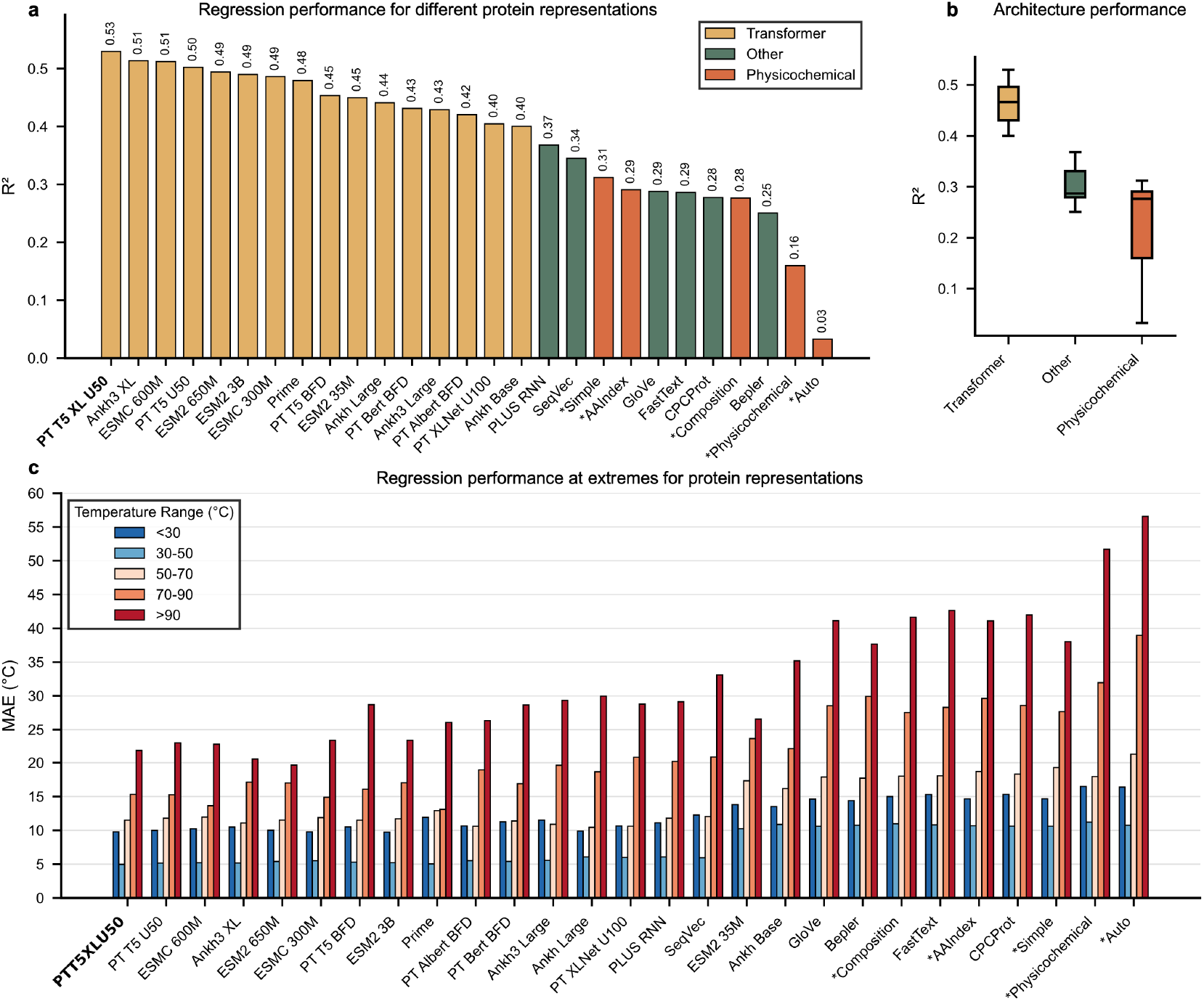
Protein representation comparison for *T*_opt_ prediction. (**a**) Coefficient of determination (*R*^2^) across protein representations, with performance averaged over Histogram Gradient Boosting, XGBoost, CatBoost, QuantileGBR, ElasticNet, SVR and KernelRidge. All models were optimised with optuna. (**b**) Performance grouped by representation architecture, comparing transformer-based protein language models (PLMs), other learned embeddings, and classical physicochemical descriptors. (**c**) Regression performance stratified by temperature range, showing that representation differences are most pronounced at temperature extremes.

Furthermore, differences in performance between protein representations are greater for extreme conditions (Fig. 4c). Only PLM embeddings maintained acceptable prediction performance, whereas simpler physicochemical descriptors and older learned representations largely failed to identify enzymes adapted to extreme conditions (i.e low predictive performance at extreme temperature ranges). Notice that PRIME, a domain adapted language model for temperature showed particularly strong performance in the 70-90°C range, while ESM-2 based representations performed best for the most extreme temperature range (>90°C), suggesting that different PLMs may capture complementary signals relevant to different temperature regions (Supplementary Table S2).

### 2.3 Resampling improves performance at extremes with minor overall prediction degradation

To address the strong imbalance in the *T*_opt_ distribution and the resulting degradation of performance at the tails, we systematically evaluated a range of resampling strategies in combination with multiple regression models. Specifically, we benchmarked both over and under-sampling approaches using CatBoost, ElasticNet, Histogram Gradient Boosting, Kernel Ridge, QuantileGBR, support vector regression (SVR), and XGBoost.

We considered several classes of resampling implemen-tations, including regression adapted algorithms such as SMOTER and WERCS, as well as “extreme” variants designed to focus explicitly on the tails of the temperature distribution. In these extreme protocols, undersampling was restricted to the central range (25-60°C) to create a more balanced training set by reducing the prevalence of mesophilic observations. Conversely, oversampling was applied only to the extremes (*<* 25, ° C and *>* 60, ° C) to generate additional tail examples. For all resampling methods hyperparameters were tuned, while keeping the downstream learning algorithms fixed at sensible hyperparameter settings to limit overall search complexity.

Across models, the strongest resampling methods yielded broadly similar outcomes: error at the extremes decreased, while MAE increased slightly for values near the median (Fig. 5). Notably, random oversampling and random undersampling targeted to the extremes performed close to the best performing algorithms, including extreme IHT and extreme ENN (Fig. 5). These latter methods are undersampling approaches that explicitly filter out observations that are difficult to predict, which may correspond to noisy or ambiguous data points. The proximity in performance suggests that, in this setting, much of the benefits comes from rebalancing exposure to tail examples rather than advanced observation selection.

**Figure 5.**
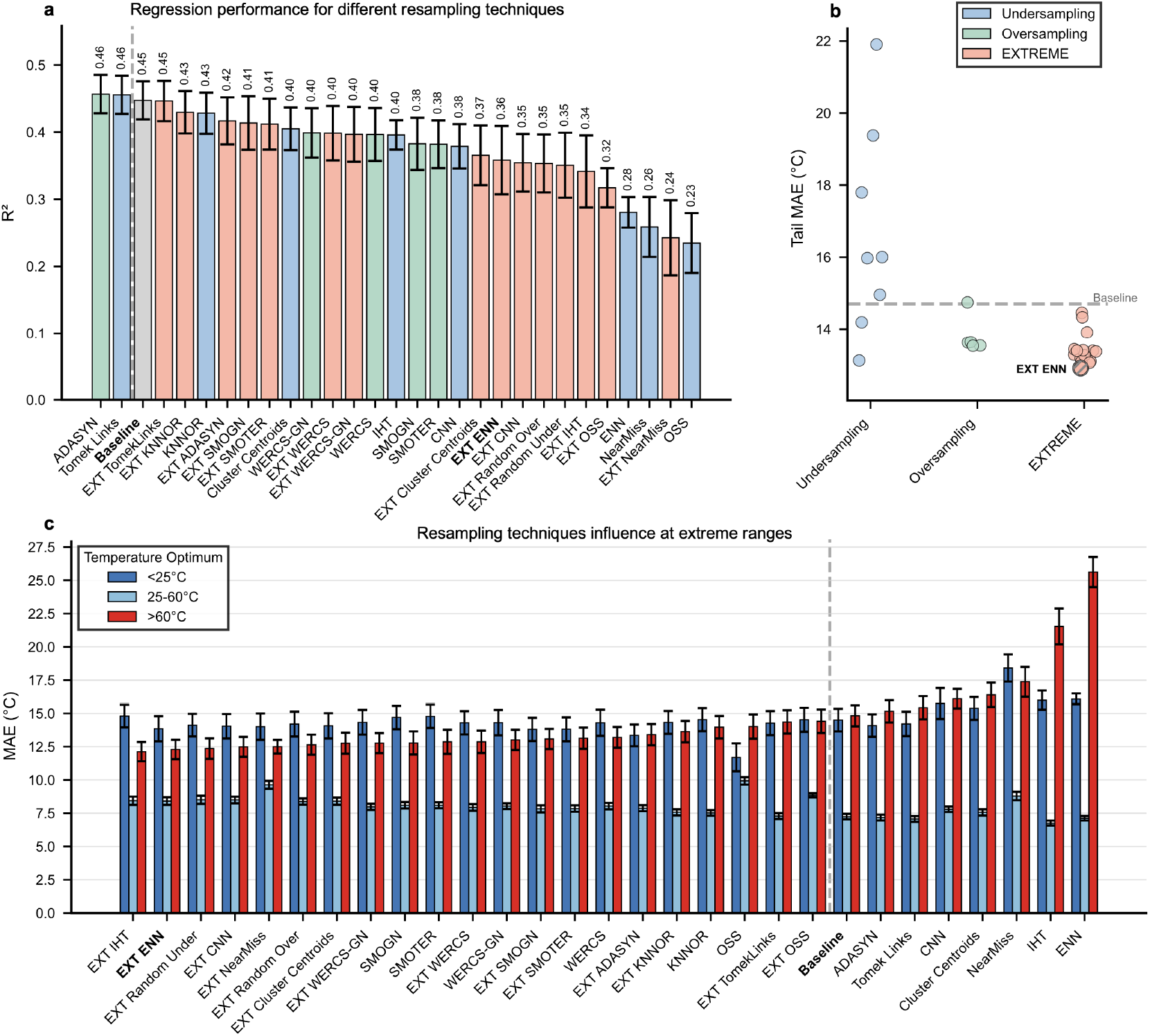
Resampling strategies for *T*_opt_ prediction. Models were trained using 5-fold cross-validation without sequence similarity correction. Resampling hyperparameters were optimized while keeping downstream regressors (Histogram Gradient Boosting, XGBoost, CatBoost, QuantileGBR, ElasticNet, SVR, and Kernel Ridge) at fixed settings. Extreme-focused implementations use masking to exclude values between 25–60 ° C for oversampling and exclude values *<* 25 °C and *>* 60 °C for undersampling. (**a**) Overall *R*^2^ comparison across resampling strategies on the held-out evaluation set. (**b**) Mean absolute error (MAE) difference relative to baseline, grouped by resampling strategy type (oversampling, undersampling, extreme-focused, and combined approaches). (**c**) MAE stratified by temperature range.

Among oversampling techniques, SMOTER and SMOGN improved tail performance relative to the baseline, but they generally remained slightly behind the best undersampling approaches. Overall, differences between resampling algorithms were modest, and only limited improvement at extremes could be achieved, indicating that resampling alone cannot fully resolve the intrinsic difficulty of extrapolating to rare temperature ranges (Supplementary Table S3).

Finally, we observed that the native IHT and ENN variants consistently worsened performance at the extremes. A plausible explanation is that, given the scarcity of extreme observations, these methods may treat tail points that lie close to mesophilic samples as hard to predict outliers and therefore remove them during instance selection. This behavior would further reduce already limited tail coverage and degrade the model’s ability to recognise enzymes adapted to extreme conditions.

Extreme-focused resampling produced clear tail gains but introduced a trade-off with global accuracy (Fig. 5b). Extreme ENN achieved the strongest tail improvement (tail MAE = 12.90 °C, Δ = −1.80 °C vs baseline) but reduced overall fit (*R*^2^ = 0.358, MAE = 9.19 °C; Fig. 5A). In contrast, extreme SMOTER and extreme WERCS provided supplementary extreme observations, reducing tail MAE to ≈ 13.40–13.43 °C with smaller degradation in overall performance (Fig. 5a–b).

### 2.4 Model comparison

We next compared regression algorithms to determine which methods were most effective for temperature prediction at extremes (Fig. 6). Overall performance was highest for ensemble and bagging-based methods, with the top models reaching *R*^2^ ≈ 0.45 (Fig. 6). Both BaggingSVR and CatBoost achieved the best overall performance for predictions at extremes.

**Figure 6.**
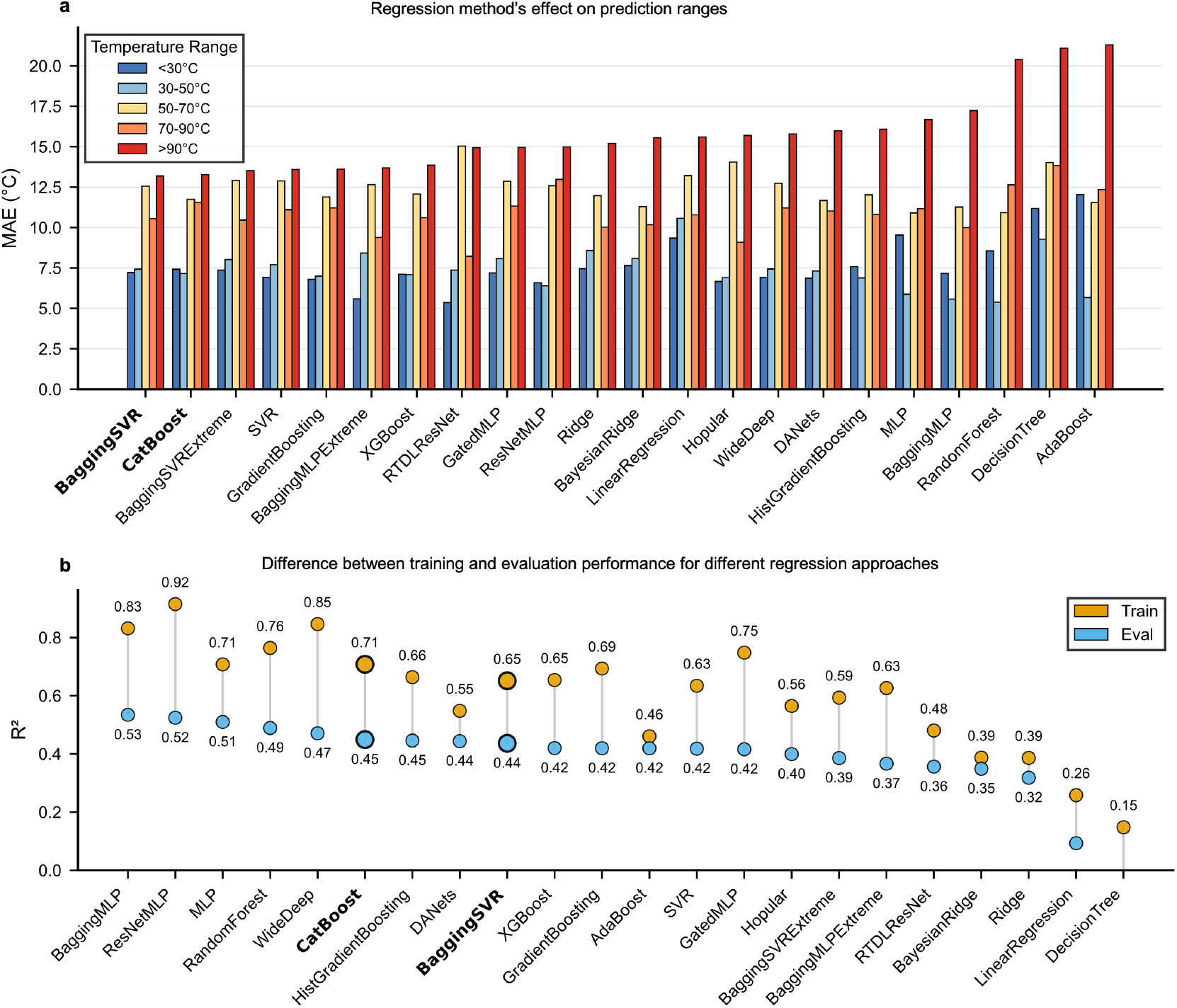
Regression model comparison for *T*_opt_ prediction. (**a**) Prediction performance across regression architectures stratified by temperature range, showing prediction error increases at temperature extremes for all models. (**b**) Difference between training and evaluation performance.

Notice that depending on the metric, many different architectures or models could be chosen. An example is GPR or KernelRdige, achieving *R*^2^ ≈ 0.56 on the evaluation data (Supplementary Table S4). However, even with regularisation, it is clear that these models fail to capture trends given their large difference between training and evaluation performance. Furthermore, other models like ResNetMLP or MLP have both high *R*^2^ values and relatively low gap between training and test performance, but fail to generalise at extreme temperatures. Improvements in global metrics did not consistently translate to improved prediction at extremes (Fig. 6C).

Tail focused analyses showed clear trade-offs between global fit and predictions at extremes. Some methods (e.g., CatBoost) showed slightly worse overall MAE on the order of ~ 1, °C relative to the best aggregate models, yet performed comparatively better at the extremes. Conversely, several ensemble and tree-based models that ranked highly by overall *R*^2^ exhibited weaker performance at extremes, and showed worse generalisation, even when measures were taken to mitigate overtraining.

Considering both overall accuracy, tail behavior, and generalisation stability (as reflected by smaller train–evaluation differences), CatBoost emerged as a reliable choice for further exploration. Finally, prediction error increased with temperature range, with values *>* 90, ° C remaining the most challenging for all models (Fig. 6A).

### 2.5 Comparison to enzyme temperature prediction models

Performance was benchmarked against the three state-of-the-art (SOTA) temperature prediction baselines using both the released models, and fully retrained models if possible (Fig 7a). For AdventML, top three major approaches were chosen for this final benchmark, training CatBoost with sigmoid importance weighting (AdventML), training Catboost with sigmoid importance weighting and with extreme ENN undersampling (AdventML tail), and a final approach adding extreme WERCS oversampling in addition to extreme ENN (AdventML extreme).

**Figure 7.**
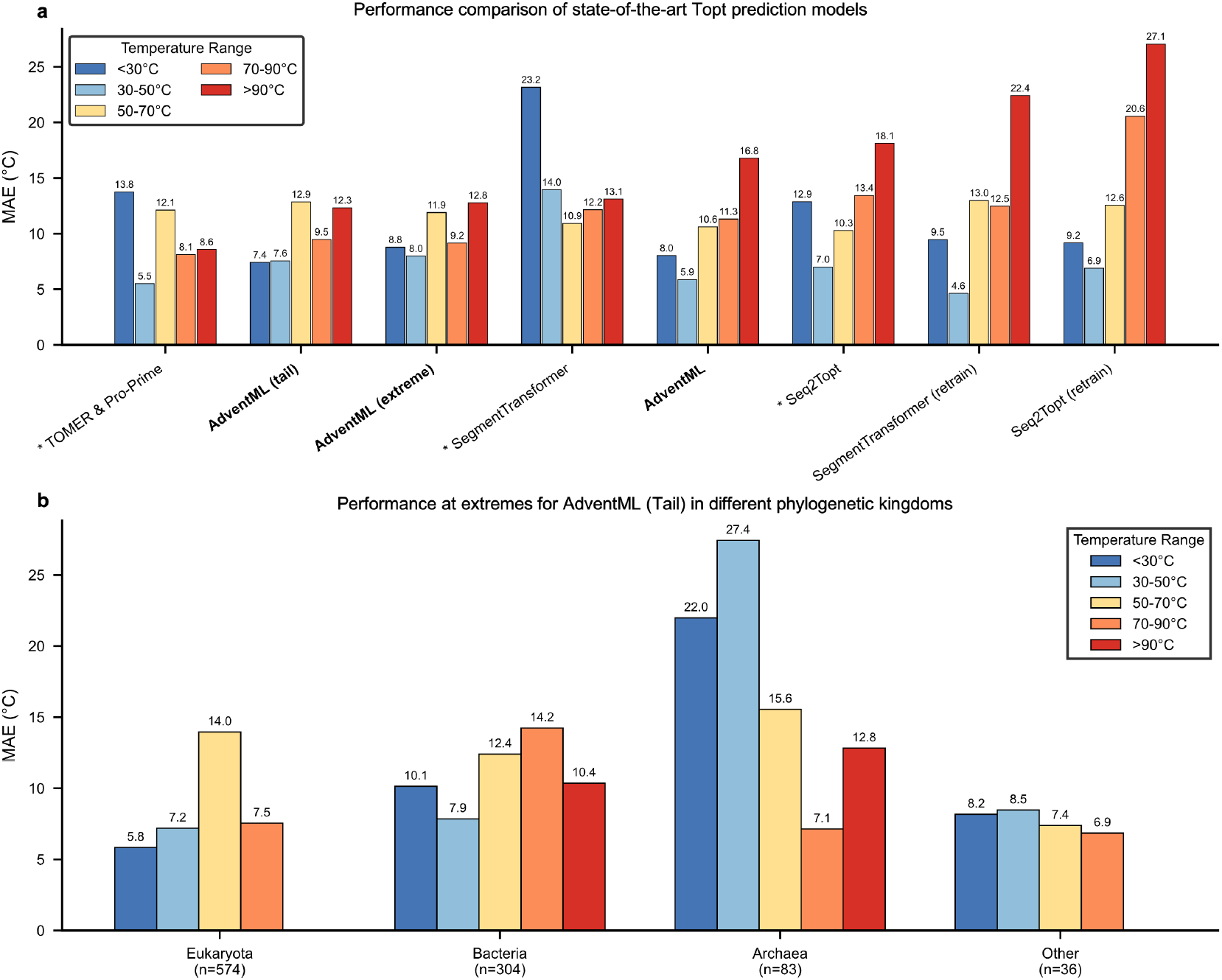
Comparison to state-of-the-art *T*_opt_ prediction models. (**a**) Performance comparison of AdventML variants against pretrained and retrained state-of-the-art models on the held-out test set. Pretrained models SegmentTransformer, TOMER and Seq2Topt may have seen part of the test data during original training, and are therefore not fully independent. AdventML models use CatBoost with Optuna hyperparameter optimization: the base variant uses no resampling, the tail variant applies extreme ENN undersampling, and the extreme variant combines extreme ENN with extreme WERCS for joint undersampling and oversampling. * indicates the models that have seen test data during training. (**b**) AdventML tail variant performance stratified by taxonomic kingdom. Limited observations in the held-out test set prevented evaluation of extreme temperature enzymes for Eukaryota and Viruses.

Overall best performance was achieved by AdventML with *R*^2^ = 0.646 and MAE = 7.58 °C (Fig. 8). However, using a more balanced metrics and taking into account prediction at extremes (Fig. 7) top performing models are AdventML tail (*R*^2^ = 0.558, MAE = 8.45 °C), AdventML extreme (*R*^2^ = 0.536, MAE = 8.85 °C) and the state-of-the-art combination of TOMER & Pro-Prime (*R*^2^ = 0.605, MAE = 8.28 °C). TOMER with Pro-Prime slightly outper-forms both AdventML tail and extreme at identifying temperature resistant enzymes (MAE_*>*90_ = 8.58 °C vs. 12.33 °C and 12.77 °C, respectively), but fails to accurately predict enzymes at lower temperatures (MAE_*<*30_ = 13.75 °C vs. 7.42 °C for AdventML tail).

**Figure 8.**
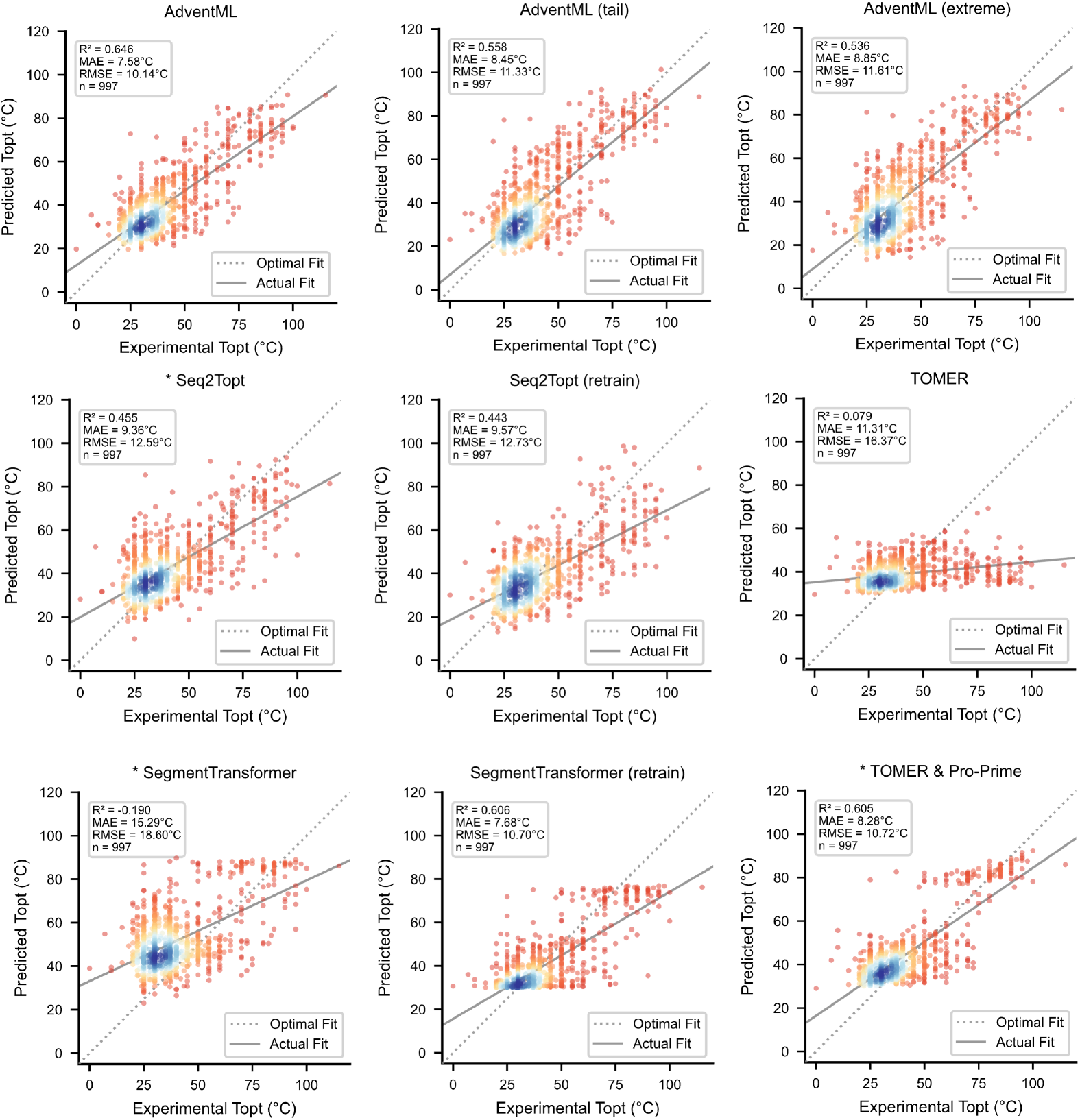
Predicted vs observed *T*_opt_ across all models. Scatter plot comparison of AdventML variants and state-of-the-art models on the held-out test set. Each panel shows predicted against observed *T*_opt_ values, with the diagonal representing perfect prediction. AdventML models use CatBoost with Optuna hyperparameter optimization: the base variant uses no resampling, the tail variant applies extreme ENN undersampling, and the extreme variant combines extreme ENN with extreme WERCS for joint undersampling and oversampling. Pretrained state-of-the-art models may have seen part of the test data during original training.

Overall, all AdventML model approaches outperform the state-of-the-art models Segment Transformer (*R*^2^ = 0.606, MAE = 7.68 °C, retrained) and Seq2Topt (*R*^2^ = 0.455, MAE = 9.36 °C). Surprisingly, only the TOMER & Pro-Prime combination outperformed our approach slightly at higher temperatures (Supplementary Table S5). These results were expected for all SOTA models, since they have seen part of the held-out test data as training input during development. This effect is largest for the TOMER Pro-Prime combination, as this approach combines PLM specific temperature embeddings trained on Brenda *T*_opt_ data (Pro-Prime), which are then used as the OGT metric in the TOMER model, effectively trained twice on parts of the held-out test set. Given the model releases on GitHub, it was not possible to retrain every model from scratch to have a completely fair compari-son without any overlap between SOTA training data and our held-out test set. Both the retrained Segment Transformer, TOMER, and TOMER & Pro-Prime combination showed a bias to overpredict temperatures (Supplementary Fig. S1). Notice the shift in performance between original model weights for SOTA models, and the retrained variants (Fig. 7). Each retrained SOTA model performed worse than its original model, an expected shift due to the complete independence of training and the held-out test set, while the original models had parts of the held-out test set as training input, leading to higher performance. For Seq2Topt, retraining shifted *R*^2^ from 0.455 to 0.443 and MAE_*>*90_ degraded sharply from 18.12 to 27.05 °C. For Segment Transformer, the original model (*R*^2^ = −0.190) improved dramatically to *R*^2^ = 0.606 after retraining, indicating the released weights were not suited to this test distribution. TOMER without OGT performed poorly (*R*^2^ = 0.079, MAE = 11.31 °C), confirming strong OGT dependence.

Differences between AdventML tail and extreme were sig-nificantly better on high temperatures than retrained SOTA models, while performance of the original Segment Transformer remained on par (Supplementary Table S6). For temperatures above 90 °C (*n* = 25) there was no significant difference between TOMER & Pro-Prime, AdventML tail/extreme, and Segment Transformer despite the large difference in Figure 7 (see also Supplementary Table S7). This was, however, expected due to the small sample size at 90 °C. Furthermore, only the TOMER & Pro-Prime combination was significantly better than our AdventML tail approach (MAE_*>*90_ = 8.58 vs. 12.33 °C), but was significantly worse at predicting cold-adapted enzymes than AdventML tail (MAE_*<*30_ = 13.75 vs. 7.42 °C; Supplementary Table S8) and has already seen testing data twice during training. For the test *R*^2^ the AdventML approach was significantly better than all other approaches (Supplementary Table S9).

### 2.6 Performance in relation to sequence and phylogenetic similarity

The dataset spans broad phylogenetic diversity (Fig. 2a), and low-dimensional structure reveals clear taxonomic organization: samples cluster by kingdom in protein feature space (Fig. 2c), while *T*_opt_ exhibits a smooth gradient across this manifold (Fig. 2d). These patterns suggest that both phylogeny and sampling density may influence generalization.

To quantify this effect directly, model performance stratified by phylogeny and by train–test similarity have been compared (Fig. S2; Table S3). Based on the simple metric of the lowest common ancestor (branch counting; range 1–14 ranks, mean = 3.1), no clear trend was observed in any model. Pearson correlations between phylogenetic distance and residuals were negligible for all AdventML variants (|*r*| *≤* 0.064) and all retrained SOTA models (|*r*| *≤* 0.055). The largest correlations occurred for TOMER (*r* = 0.180), Segment Transformer (*r* = 0.093) and Seq2Topt (*r* = 0.066), but they still showed barely any correlation. Furthermore, no deviation could be observed for models based on sequence similarity. Higher sequence similarity did not result in better predictions or a clear trend, with Pearson correlations between similarity and absolute prediction error near zero for all models (|*r*| *≤* 0.068). Only deviations can be observed for Segment Transformer with a tendency to predict temperatures higher than the actual experimental values (MBE = 11.75 °C), and a similar trend in the TOMER & Pro-Prime combination, but to a lesser degree (MBE = 3.72 °C).

## 3. Discussion

This work investigated whether enzyme optimal temperature prediction can be improved by combining protein language model (PLM) embeddings with resampling strategies and machine learning architecture search. The results demonstrate that PLMs outperform classical biology based features. Furthermore, resampling improved performance at extremes, with simple approaches performing better or on par with more complex algorithms. Finally, our best model combined catboost with extreme ENN undersampling to improve predictions for Topt significantly compared to SOTA models.

The results demonstrate that PLM embeddings substantially outperform classical sequence-derived features for *T*_opt_ prediction. However, among the top-performing PLMs, differences were small, and neither model size nor publication date showed a consistent trend. For example, ESM-2 650M and 3B performed comparably, and newer models such as ESMC 300M did not outperform older models like ProtT5 XL U50. This indicates that for state-of-the-art embeddings, practical considerations such as licensing and inference speed may be used as additional selection criteria, rather than parameter count or the most recent model. A similar observation was made in literature, were smaller older models can outperform recent ones, especially on problems with limited data^29,30^. Incorporating OGT as an auxiliary feature provided clear gains for classical repre-sentations, but only marginal improvements when combined with PLMs, indicating that PLMs already capture much of the organism level thermal information.

Resampling consistently improved prediction at temperature extremes at the cost of a minor overall R! penalty, in line with findings reported for TOMER^10^. The present study extended this analysis to a broader set of undersampling and hybrid over/undersampling methods, including WERCS and SMOGN. Even after hyperparameter optimisation, targeted random undersampling in specific temperature ranges matched or outperformed these more complex approaches, indicating that complex resampling algorithms do not necessarily translate to prediction gains. Notably, resampling provided improvements beyond corrections such as weighted loss functions, underscoring its importance for rare event prediction^31^.

Across machine learning types, large overall performance differences were observed, but variation among top-performing methods was minor. The final model, ProtT5 XL U50 combined with CatBoost and Extreme ENN under-sampling, achieved performance exceeding current state-of-the-art models. CatBoost was selected over BaggingSVR despite negligible performance differences, given its lower computational cost and greater interpretability. Advanced deep learning architectures such as Gated MLP and ResNetMLP did not improve over gradient-boosted or kernel-based methods, likely due to the low observation-to-feature ratio limiting their generalisation capacity. Furthermore, analyses stratified by phylogenetic distance and sequence similarity revealed no clear trend of prediction bias toward closer evolutionary relatedness, suggesting that the model generalises robustly across the taxonomic range present in the data.

Current prediction performance appears to plateau around 0.5-0.6 *R*^2^, and it remains unclear whether substantial further gains are achievable given inherent limitations in the underlying data. *T*_opt_ values, even when sourced from a single database such as BRENDA, are notoriously noisy: duplicate entries for the same enzyme frequently report widely different values, and key experimental variables such as buffer composition, substrate identity, pH, and assay or experimental duration, are often absent for predictions. As a result, predicting *T*_opt_ from protein sequence or structure alone is inherently an approximation of a context-dependent target variable. This challenge is further increased by the nature of thermal activity profiles, where enzymes can lack a sharp optimum and instead exhibit broad activity plateaus across a range of temperatures^32^. Such profiles introduce substantial variability in both training labels and evaluation targets, effectively imposing a sort of ceiling on achievable prediction accuracy that reflects experimental noise rather than model deficiency.

Limitations of the AdventML approach to selecting an optimal prediction pipeline is the separation of embeddings, resampling and algorithm testing. Ideally, resampling approaches should be tested together with different machine learning algorithms, since these can influence the outcome^33^. Additionally, a fixed training-validation-test ap-proach was chosen for selection since this allowed a fair comparisons taking sequence similarity into account. Ideally, such an approach can be supplemented with bootstrapping or Monte-Carlo cross-fold validation to gain insights in the model variability. Further expansion and improvement of *T*_opt_ is possible with some interesting approaches. Firstly, while in AdventML an optimal PLM was chosen, a combination of several distinct PLMs might enrich feature space. Secondly, given the current advances in models, approaches using pseudo-labels from existing models such as AdventML, Segment Transformer or Seq2Topt might be interesting to try and enrich models further. And while finetuning has been done in Seq2Topt’s approach^34^, using related information such as protein thermostability might further improve this type of models.

## Conclusion

AdventML is a prediction framework for enzyme optimal catalytic temperature that combines protein language model embeddings with resampling strategies and classical machine learning. The results demonstrate that modern PLM embeddings render organism growth temperature largely redundant for *T*_opt_ prediction, and that extreme-focused re-sampling improves tail predictions at minimal cost to overall accuracy. Notably, CatBoost with ProtT5 XL U50 embeddings and extreme ENN undersampling outperformed current state-of-the-art models, including deep learning approaches specifically designed for temperature prediction. AdventML is freely available as a GitHub tool to support the identification of thermostable and cold-adapted enzymes for biotechnological applications.

## 4. Methods

### 4.1 Workflow overview

A pipeline was developed to predict the enzyme optimal temperature (*T*_opt_) from protein sequences by examining different prediction strategies.

First, *T*_opt_ and sequence data were extracted from the BRENDA enzyme database and preprocessed, including quality filtering, deduplication, and taxonomic annotation. To prevent information leakage, sequences were clustered by identity and separated into a training set and a held-out test set, with both sequence similarity and organism-level constraints.

Second, a systematic comparison was performed across multiple protein representations (transformer-based embeddings, other learned embeddings, and classical physico-chemical descriptors), resampling strategies for addressing temperature distribution imbalance, and regression architectures. Hyperparameters were tuned via Bayesian optimization, and all models were evaluated using standardized metrics across temperature ranges.

Finally, an optimal temperature prediction pipeline configuration was selected, final models were trained, and performance was benchmarked against existing state-of-the-art *T*_opt_ prediction methods on the held-out test set, independent from earlier architectural comparisons.

### 4.2 Data collection and preprocessing

The optimal temperature (*T*_opt_) dataset was obtained from the BRENDA enzyme database^35^ through extraction from the BRENDA JSON file (version 2025_01) with release date May 30th 2025. For each entry, *T*_opt_ values were retrieved together with organism name, UniProt accession, and experimental comments. Next, Taxonomy IDs were assigned using the ETE3 NCBITaxa toolkit^36^ based on organism names. UniProt sequences were retrieved from UniProt^37^ via the UniProt REST API using accession IDs; obsolete accessions were remapped if possible. Entries without UniProt IDs from BRENDA were matched with the EC number and organism ID, and flagged accordingly.

The raw dataset underwent quality control adapted from^9^. Sequences with fewer than 50 amino acids or more than 4000 amino acids were excluded. Entries where *T*_opt_ val-ues were annotated with “assay at” in the BRENDA comments field were removed to ensure true optimal temperatures rather than arbitrary assay conditions. Duplicate entries sharing UniProt identifiers, identical *T*_opt_ values, and matching EC number prefixes were merged. For enzymes with multiple *T*_opt_ values, mean and standard deviation were calculated; outlier values deviating by more than *ω* 10 °C from the mean were excluded before computing the average. Only true *T*_opt_ not originating from average calculations or without duplicates were considered for a held-out test set. To ensure robust evaluation and prevent data leakage, sequences were partitioned into training and test sets using sequence similarity clustering with MMseqs2^38^. Sequences were clustered at 90% identity with a bidirectional 80% coverage constraint to ensure only true full-length homologs were grouped together, meaning that test sequences shared less than 90% identity with any training sequence. Additionally, organism-level constraints were added such that no species appeared in both training and test sets. The heldout test set contained 10% of the original dataset, with the remaining 90% used for training. The test set (held-out set) was only used in the final evaluation comparing state-of-the-art models with full pipeline implementations, other analysis such as protein representation comparison or resampling methods comparison were done with only the training set. For OGT prediction experiments, we utilized the dataset from^10^, containing 2,917 organism temperature pairs. The original split from Tomer et al. was used for training and testing. Furthermore, the dataset was used only to make a comparison between predictions with and without OGT for different protein representations.

### 4.3 Protein features and representations

Protein sequences were encoded using multiple protein representations to examine the optimal feature space for *T*_opt_ prediction. Three major protein representation groups were examined, including transformer-based embeddings, other learned protein representations and biology or physicochemical derived protein features. For transformer-based embeddings, average pooling was used to create fixed-length sequence-level embeddings. We examined ESM-2 (650M and 3B parameter variants)^39^, ESM-C (600M), ESM-1^40^, PRIME, xTrimoPGLM, ProtTrans models (T5-XL-U50, T5-UniRef50, BERT, ALBERT, XLNet)^41^, and Ankh variants^42^. Other learned protein representations we used include SeqVec^43^, UniRep^44^, PlusRNN^45^, CPCProt^46^, Bepler, FastText^47^, GloVe^48^, Word2Vec^49^. Furthermore, we calculated physicochemical properties using protlearn, including amino acid composition, pseudo-amino acid composition (PAAC), conjoint triad descriptors (CTD), autocorrelation descriptors, and AAindex-derived properties. These interpretable baselines captured basic compositional biases without learned representations. All feature representations are listed in Supplementary Table S10.

### 4.4 Protein embeddings comparison

Training data was split with MMseqs2 at 90% sequence identity into training, validation, and evaluation sets (70%/15%/15%). Each representation was evaluated across seven regression models: HistGradientBoosting, XGBoost, CatBoost, Quantile Gradient Boosting, ElasticNet, Support Vector Regression, and Kernel Ridge Regression. Hyperparameters were optimized using Optuna^50^ with 96 trials, minimizing a two-tailed sigmoid-weighted MAE with hard cutoffs at 25 °C and 60 °C to emphasize performance at temperature extremes. All hyperparameters were selected based on validation set performance. Models were evaluated on the held-out evaluation set, and average performance across all regressors was computed to provide a representation-level comparison independent of model choice.

### 4.5 *T*_opt_ prediction in the presence or absence of organism growth temperature

The influence of OGT on *T*_opt_ prediction was assessed for different feature sets. Data with annotated sequence, *T*_opt_, and OGT values (derived from the TOMER dataset) were used, keeping the original splits from Gado et al.^10^ Three regressors (HistGradientBoosting, CatBoost, Support Vector Regression) were trained in two modes: with OGT included as a feature and without OGT. Hyperparameter optimization with 96 trials followed the same sigmoid-weighted MAE objective. For each representation, performance was averaged across the three models to isolate the contribution of OGT.

### 4.6 Resampling strategies

Enzyme *T*_opt_ values exhibited imbalance, with heavy concentration in the mesophilic range (30–50 °C) and sparse representation at low (*<* 25 °C) and high (*>* 60 °C) extremes. To improve model sensitivity across the full temperature spectrum, a selection of resampling algorithms for continuous data was evaluated on the training set only.

#### Undersampling methods

Condensed Nearest Neigh-bours (CNN), Cluster Centroids, Edited Nearest Neighbours (ENN), Instance Hardness Threshold (IHT), NearMiss, One-Sided Selection (OSS), and Tomek Links.

#### Oversampling methods

SMOTER (SMOTE for regression), SMOGN (SMOTE with Gaussian noise), ADASYN, *k*-NN oversampling (KNNOR), and WERCS (weighted relevance-based combination strategy).

#### Extreme-focused variants

Tail-specific versions of the above methods (e.g., extreme ENN, extreme SMOTER) that selectively oversample the rare low and high temperature domains (*<* 25 °C and *>* 60 °C) while undersampling the dense mid-range (25-60 °C).

For 5-fold cross validation, per fold, features were standardized, the resampling strategy was applied, and seven regressors were trained. The trained regressors were Histogram Gradient Boosting, XGBoost, CatBoost, QuantileGBR, ElasticNet, SVR and KernelRidge. For each resampling strategy, several resampling hyperparameters were tried with a grid-search approach, keeping regression hyperparameters fixed at a sensible setting. Performance per fold was averaged over the seven regression models. All resampling approaches are listed in Supplementary Table S11 and S12.

### 4.7 Model comparison and evaluation

To evaluate *T*_opt_ prediction performance on ProtTrans T5-XL-U50 embeddings, a set of classical regression and deep learning architectures was benchmarked.

The regression cohort consisted of 29 individual model from 5 model families. **Linear models:** Lasso, Ridge, Elastic-Net, Bayesian Ridge, Huber, Linear Regression, Stochastic Gradient Descent (SGD), Passive Aggressive, Quantile Regression, Orthogonal Matching Pursuit (OMP), Automatic Relevance Determination (ARD), and Partial Least Squares (PLS). **Tree-based models:** Decision Tree, Random Forest, Extra Trees, Gradient Boosting, HistGradientBoosting, and AdaBoost. **Gradient boosting libraries:** XGBoost, Light-GBM, and CatBoost. **Distance- and kernel-based models:** *k*-Nearest Neighbours (KNN), Support Vector Regression (SVR), Kernel Ridge, and Gaussian Process Regression (GPR). **Neural:** Multi-Layer Perceptron (MLP). In addition, weighted bagging ensembles were constructed for four base learners (CatBoost, MLP, SVR, Decision Tree), each tested at two oversampling intensities (moderate 10 × and extreme 1000 ×), resulting in eight additional ensemble configurations.

Furthermore, 22 tabular deep learning architectures were evaluated: Baseline MLP, RTDL ResNet, RealMLP, TabM, Gated MLP, Sparse MLP, TabNet, LassoNet, GAN-DALF, DANets, TabR, Hopular, FT-Transformer, SAINT, NODE, GrowNet, VIME, ResNet MLP, Wide&Deep, TabNet-Inspired, Attention MLP, and SNN. All deep learning architecture names are listed in Supplementary Table S13.

Models were evaluated on the training data with a 70%-15%-15% spit for training, validation and evaulation data. Hyperparameters for all architectures were optimized using Bayesian optimization via Optuna. Classical regression models used 192 TPE trials, whereas deep learning architectures utilized 100 TPE trials. Deep learning training relied on a maximum of 500 epochs with early stopping (30 epochs patience) and learning-rate reduction on plateau (15 epochs patience). Specifically, VIME employed a true pretraining phase (100 final epochs, batch size 256, mask probability 0.3) before fine-tuning, and GrowNet used stagewise boosting with 50 epochs per stage. All hyperparameters are listed in Supplementary Table S14.

To address the severe target imbalance, sample weights were applied during training and hyperparameter optimization using a two-tailed sigmoid function with exact cutoffs at 25 °C and 60 °C. This resulted in higher unit weight at temperature extremes while down weighting importance in the mesophilic core. The dynamic weight range applied was 1000× (minimum weight 0.001) for classical regression models. For deep learning architectures, this range was intentionally narrowed to 100× (minimum weight 0.01) to preserve gradient magnitudes and training stability across large parameter spaces.

Performance was quantified using standard regression metrics: coefficient of determination (*R*^2^), root-mean-square error (RMSE), mean absolute error (MAE), mean absolute percentage error (MAPE), mean bias error (MBE), and Spearman and Pearson correlation coefficients. Metrics were reported for the overall test set, isolated tail regions (*T*_opt_ *<*20 °C or *>* 60 °C), and within stratified temperature bins (*<* 30, 30-50, 50-70, 70-90, *>* 90 °C).

### 4.8 AdventML pipeline evaluation

The model CatBoost was combined with the ProtTrans T5 XL UniRef50 embeddings and tested with extreme ENN and extreme ENN/WERCS for the development of the final *T*_opt_ prediction model. Models AdventML, AdventML tail and AdventML extreme were developed, using catboost without resampling, with extreme ENN resampling, and with the combination of extreme ENN and extreme WERCS resampling. Hyperparameters were optimized using Optuna with 192 TPE trials, maximizing weighted *R*^2^ on the validation set (15% of training data). Final models were retrained on the full resampled training data with optimized hyperparameters and sigmoid sample weighting (cutoffs 2560 °C, dynamic range 1000*×*), then evaluated on the held-out test set.

### 4.9 Comparison to state-of-the-art

External baseline methods were retrained and evaluated on the same leakage-safe data splits to enable direct comparison. These included:

- **Seq2Topt**: ESM-2 encoder (esm2_t6_8M_UR50D, 8M parameters) with multi-head attention and residual dense blocks, trained with MSE loss and Adam optimizer, using min–max target scaling to the 0–120 °C range^34^.
- **TOMER**: Rebagg ensemble of 100 decision tree regressors using amino acid composition (20 features) and OGT (1 feature) as input, with sigmoid-relevance resampling. Evaluated both with annotated OGT and with a default OGT of 37 °C when OGT was unavailable^10^.
- **SegmentTransformer**: Multi-scale segment attention network with a frozen ESM-2 (150M) backbone. Sequences truncated to 1000 residues^26^.
- **PRIME**: Pre-trained protein language model (AI4Protein/Prime_690M) used as an OGT predictor. Predicted OGT values were supplied as input features to TOMER when experimental OGT was unavailable^28^.

Each baseline produced test set predictions aligned with the AdventML evaluation framework, enabling head-to-head comparison under identical data partitions and evaluation metrics.

### 4.10 Statistical significance testing

Statistical significance of performance differences between models was assessed using the paired bootstrap resampling test following the methodology of Koehn et al.^51,52^. All state-of-the-art models were compared, including AdventML variants and retrained baselines. For each of the unique model pairs, *B* = 100,000 bootstrap replicates were drawn by sampling *N* = 997 aligned test predictions with replacement, using identical indices for both models in each replicate to preserve the paired structure. For each bootstrap sample, the corpus-level metric was computed for both models and the number of wins for each model were recorded. Tied scores for a given bootstrap sample were counted as 0.5 for both models. Two-sided *p*-values were computed as 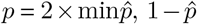, with 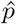 the proportion of bootstrap replicates in which model A achieved a better score than model B. To control the false discovery rate across all pairwise comparisons for each metric, *p*-values were adjusted using the Benjamini–Hochberg procedure at *α* = 0.05^53^. Comparisons were performed for *R*^2^, MAE, and MAE within temperature subsets (*<* 30 °C, *>* 70 °C, *>* 90 °C).

## Data and Code Availability

The AdventML prediction models and code are freely available on GitHub at https://github.com/LegallyOverworked/adventml. The processed *T*_opt_ dataset and trained models are available at [repository/DOI].

## Author Contributions

J.F. designed the study, developed the methodology, performed the analyzes, and wrote the manuscript. V.v.N and B.D.M. supervised the project and provided scientific feedback.

## Bibliography

1. David C Demirjian, Francisco Moris-Varas, and Constance S Cassidy. Enzymes from extremophiles. Current Opinion in Chemical Biology, 5(2):144–151, April 2001. ISSN 13675931. doi: 10.1016/S1367-5931(00)00183-6. URL https://linkinghub.elsevier.com/retrieve/pii/S1367593100001836.

2. Pernilla Turner, Gashaw Mamo, and Eva Nordberg Karlsson. Potential and utilization of thermophiles and thermostable enzymes in biorefining. Microbial Cell Factories, 6(1):9, December 2007. ISSN 1475-2859. doi: 10.1186/1475-2859-6-9. URL https://microbialcellfactories.biomedcentral.com/articles/10.1186/1475-2859-6-9.

3. Claire Vieille and Gregory J. Zeikus. Hyperthermophilic Enzymes: Sources, Uses, and Molecular Mechanisms for Thermostability. Microbiology and Molecular Biology Reviews, 65(1): 1–43, March 2001. ISSN 1092-2172, 1098-5557. doi: 10.1128/MMBR.65.1.1-43.2001. URL https://journals.asm.org/doi/10.1128/MMBR.65.1.1-43.2001.

4. G Haki. Developments in industrially important thermostable enzymes: a review. Biore-source Technology, 89(1):17–34, August 2003. ISSN 09608524. doi: 10.1016/S0960-8524(03)00033-6.pii/S0960852403000336. URL https://linkinghub.elsevier.com/retrieve/

5. Skander Elleuche, Carola Schröder, Kerstin Sahm, and Garabed Antranikian. Extremozymes—biocatalysts with unique properties from extremophilic microorganisms. Current Opinion in Biotechnology, 29:116–123, October 2014. ISSN 09581669. doi: 10.1016/j.copbio.2014.04.003. URL https://linkinghub.elsevier.com/retrieve/pii/S0958166914000755.

6. Sonia Jemli, Dorra Ayadi-Zouari, Hajer Ben Hlima, and Samir Bejar. Biocatalysts: application and engineering for industrial purposes. Critical Reviews in Biotechnology, 36(2):246– 258, March 2016. ISSN 0738-8551, 1549-7801. doi: 10.3109/07388551.2014.950550. URL http://www.tandfonline.com/doi/full/10.3109/07388551.2014.950550.

7. Carl J. Yeoman, Yejun Han, Dylan Dodd, Charles M. Schroeder, Roderick I. Mackie, and Isaac K.O. Cann. Thermostable Enzymes as Biocatalysts in the Biofuel Industry. In Advances in Applied Microbiology, volume 70, pages 1–55. Elsevier, 2010. ISBN 978-0-12-380991-9. doi: 10.1016/S0065-2164(10)70001-0. URL https://linkinghub.elsevier.com/retrieve/pii/S0065216410700010.

8. Sumit Kumar, Arun K. Dangi, Pratyoosh Shukla, Debabrat Baishya, and Sunil K. Khare. Thermozymes: Adaptive strategies and tools for their biotechnological applications. Bioresource Technology, 278:372–382, April 2019. ISSN 09608524. doi: 10.1016/j.biortech.2019.01.088. URL https://linkinghub.elsevier.com/retrieve/pii/S0960852419301087.

9. Gang Li, Kersten S. Rabe, Jens Nielsen, and Martin K. M. Engqvist. Machine Learning Applied to Predicting Microorganism Growth Temperatures and Enzyme Catalytic Optima. ACS Synthetic Biology, 8(6):1411–1420, June 2019. ISSN 2161-5063, 2161-5063. doi: 10.1021/acssynbio.9b00099. URL https://pubs.acs.org/doi/10.1021/acssynbio.9b00099. TOME.

10. Japheth E. Gado, Gregg T. Beckham, and Christina M. Payne. Improving Enzyme Optimum Temperature Prediction with Resampling Strategies and Ensemble Learning. Journal of Chemical Information and Modeling, 60(8):4098–4107, August 2020. ISSN 1549-9596, 1549-960X. doi: 10.1021/acs.jcim.0c00489. URL https://pubs.acs.org/doi/10.1021/acs.jcim.0c00489. TOMER.

11. Martin K. M. Engqvist. Correlating enzyme annotations with a large set of microbial growth temperatures reveals metabolic adaptations to growth at diverse temperatures. BMC Microbiology, 18(1):177, December 2018. ISSN 1471-2180. doi: 10.1186/s12866-018-1320-7. URL https://bmcmicrobiol.biomedcentral.com/articles/10.1186/s12866-018-1320-7.

12. Michael S. Rappé and Stephen J. Giovannoni. The Uncultured Microbial Majority. Annual Review of Microbiology, 57(1):369–394, October 2003. ISSN 0066-4227, 1545-3251. doi: 10.1146/annurev.micro.57.030502.090759. URL https://www.annualreviews.org/doi/10.1146/annurev.micro.57.030502.090759.

13. Antje Chang, Lisa Jeske, Sandra Ulbrich, Julia Hofmann, Julia Koblitz, Ida Schomburg, Meina Neumann-Schaal, Dieter Jahn, and Dietmar Schomburg. BRENDA, the ELIXIR core data resource in 2021: new developments and updates. Nucleic Acids Research, 49(D1):D498–D508, January 2021. ISSN 0305-1048, 1362-4962. doi: 10.1093/nar/gkaa1025. URL https://academic.oup.com/nar/article/49/D1/D498/5992283.

14. Donal A Hickey and Gregory Ac Singer. Genomic and proteomic adaptations to growth at high temperature. Genome Biology, 5(10):117, 2004. ISSN 14656906. doi: 10.1186/gb-2004-5-10-117. URL http://genomebiology.biomedcentral.com/articles/10.1186/gb-2004-5-10-117.

15. H. Nakashima. Compositional Changes in RNA, DNA and Proteins for Bacterial Adaptation to Higher and Lower Temperatures. Journal of Biochemistry, 133(4):507–513, April 2003. ISSN 0021924X. doi: 10.1093/jb/mvg067. URL https://academic.oup.com/jb/article-lookup/doi/10.1093/jb/mvg067.

16. Nicolas Galtier and J.R. Lobry. Relationships Between Genomic G+C Content, RNA Secondary Structures, and Optimal Growth Temperature in Prokaryotes. Journal of Molecular Evolution, 44(6):632–636, June 1997. ISSN 0022-2844, 1432-1432. doi: 10.1007/PL00006186. URL http://link.springer.com/10.1007/PL00006186.

17. Konstantin B Zeldovich, Igor N Berezovsky, and Eugene I Shakhnovich. Protein and DNA Sequence Determinants of Thermophilic Adaptation. PLoS Computational Biology, 3(1):e5, January 2007. ISSN 1553-7358. doi: 10.1371/journal.pcbi.0030005. URL https://dx.plos.org/10.1371/journal.pcbi.0030005.

18. Lucas Sawle and Kingshuk Ghosh. How Do Thermophilic Proteins and Proteomes Withstand High Temperature? Biophysical Journal, 101(1):217–227, July 2011. ISSN 00063495. doi: 10.1016/j.bpj.2011.05.059. URL https://linkinghub.elsevier.com/retrieve/pii/S0006349511006618.

19. Gregory A.C. Singer and Donal A. Hickey. Thermophilic prokaryotes have characteristic patterns of codon usage, amino acid composition and nucleotide content. Gene, 317:39–47, October 2003. ISSN 03781119. doi: 10.1016/S0378-1119(03)00660-7. URL https://linkinghub.elsevier.com/retrieve/pii/S0378111903006607.

20. M.Michael Gromiha, Motohisa Oobatake, and Akinori Sarai. Important amino acid properties for enhanced thermostability from mesophilic to thermophilic proteins. Biophysical Chemistry, 82(1):51–67, November 1999. ISSN 03014622. doi: 10.1016/S0301-4622(99)00103-9. URL https://linkinghub.elsevier.com/retrieve/pii/S0301462299001039.

21. Nan Zhao, Bin Pang, Chi-Ren Shyu, and Dmitry Korkin. Charged residues at protein interaction interfaces: Unexpected conservation and orchestrated divergence. Protein Science, 20(7):1275–1284, July 2011. ISSN 0961-8368, 1469-896X. doi: 10.1002/pro.655. URL https://onlinelibrary.wiley.com/doi/10.1002/pro.655.

22. András Szilágyi and Péter Závodszky. Structural differences between mesophilic, moderately thermophilic and extremely thermophilic protein subunits: results of a comprehensive survey. Structure, 8(5):493–504, May 2000. ISSN 09692126. doi: 10.1016/S0969-2126(00)00133-7. URL https://linkinghub.elsevier.com/retrieve/pii/S0969212600001337.

23. Jeremy L. England, Boris E. Shakhnovich, and Eugene I. Shakhnovich. Natural selection of more designable folds: A mechanism for thermophilic adaptation. Proceedings of the National Academy of Sciences, 100(15):8727–8731, July 2003. ISSN 0027-8424, 1091-6490. doi: 10.1073/pnas.1530713100. URL https://pnas.org/doi/full/10.1073/pnas.1530713100.

24. Roger L. Chang, Kathleen Andrews, Donghyuk Kim, Zhanwen Li, Adam Godzik, and Bernhard O. Palsson. Structural Systems Biology Evaluation of Metabolic Thermotolerance in Escherichia coli. Science, 340(6137):1220–1223, June 2013. ISSN 0036-8075, 1095-9203. doi: 10.1126/science.1234012. URL https://www.science.org/doi/10.1126/science.1234012.

25. Sofia Khan and Mauno Vihinen. Performance of protein stability predictors. Human Mutation, 31(6):675–684, March 2010. ISSN 10597794, 10981004. doi: 10.1002/humu.21242. URL https://onlinelibrary.wiley.com/doi/10.1002/humu.21242.

26. Ziqi Zhang, Shiheng Chen, Runze Yang, Zhisheng Wei, Wei Zhang, Lei Wang, Zhanzhi Liu, Fengshan Zhang, Jing Wu, Xiaoyong Pan, Hongbin Shen, Longbing Cao, and Zhaohong Deng. Modeling Enzyme Temperature Stability from Sequence Segment Perspective. Journal of Chemical Information and Modeling, page acs.jcim.5c01674, October 2025. ISSN 1549-9596, 1549-960X. doi: 10.1021/acs.jcim.5c01674. URL https://pubs.acs.org/doi/10.1021/acs.jcim.5c01674. Segment Transformer.

27. Sizhe Qiu, Bozhen Hu, Jing Zhao, Weiren Xu, and Aidong Yang. _eprint: a sequence-based deep learning predictor of enzyme optimal temperature. bioRxiv, 2024. doi: 10.1101/2024.08.12.607600. URL https://www.biorxiv.org/content/early/2024/08/14/2024.08.12.607600. https://www.biorxiv.org/content/early/2024/08/14/2024.08.12.607600.full.pdf.

28. Fan Jiang, Mingchen Li, Jiajun Dong, Yuanxi Yu, Xinyu Sun, Banghao Wu, Jin Huang, Liqi Kang, Yufeng Pei, Liang Zhang, Shaojie Wang, Wenxue Xu, Jingyao Xin, Wanli Ouyang, Guisheng Fan, Lirong Zheng, Yang Tan, Zhiqiang Hu, Yi Xiong, Yan Feng, Guangyu Yang, Qian Liu, Jie Song, Jia Liu, Liang Hong, and Pan Tan. A general temperature-guided language model to design proteins of enhanced stability and activity. Science Advances, 10 (48):eadr2641, November 2024. ISSN 2375-2548. doi: 10.1126/sciadv.adr2641. URL https://www.science.org/doi/10.1126/sciadv.adr2641.

29. Tobias Senoner, Ivan Koludarov, Joshua Günther, Amarda Shehu, Burkhard Rost, and Yana Bromberg. Which pLM to choose?, October 2025. URL http://biorxiv.org/lookup/doi/10.1101/2025.10.30.685515.

30. Luiz C. Vieira, Morgan L. Handojo, and Claus O. Wilke. Medium-sized protein language models perform well at transfer learning on realistic datasets. Scientific Reports, 15(1):21400, July 2025. ISSN 2045-2322. doi: 10.1038/s41598-025-05674-x. URL https://www.nature.com/articles/s41598-025-05674-x.

31. Yuzhe Yang, Kaiwen Zha, Ying-Cong Chen, Hao Wang, and Dina Katabi. Delving into Deep Imbalanced Regression, 2021. URL https://arxiv.org/abs/2102.09554. Version Number: 2.

32. Vitor M. Almeida and Sandro R. Marana. Optimum temperature may be a misleading parameter in enzyme characterization and application. PLOS ONE, 14(2):e0212977, February 2019. ISSN 1932-6203. doi: 10.1371/journal.pone.0212977. URL https://dx.plos.org/10.1371/journal.pone.0212977.

33. Juscimara G. Avelino, George D. C. Cavalcanti, and Rafael M. O. Cruz. Resampling strategies for imbalanced regression: a survey and empirical analysis. Artificial Intelligence Review, 57 (4):82, March 2024. ISSN 1573-7462. doi: 10.1007/s10462-024-10724-3. URL https://link.springer.com/10.1007/s10462-024-10724-3.

34. Sizhe Qiu, Bozhen Hu, Jing Zhao, Weiren Xu, and Aidong Yang. Seq2Topt: a sequence-based deep learning predictor of enzyme optimal temperature. Briefings in Bioinformatics, 26(2):bbaf114, March 2025. ISSN 1467-5463, 1477-4054. doi: 10.1093/bib/bbaf114. URL https://academic.oup.com/bib/article/doi/10.1093/bib/bbaf114/8074761. Seq2Topt.

35. Ida Schomburg, Antje Chang, Oliver Hofmann, Christian Ebeling, Frank Ehrentreich, and Dietmar Schomburg. BRENDA: a resource for enzyme data and metabolic information. Trends in Biochemical Sciences, 27(1):54–56, January 2002. ISSN 09680004. doi: 10.1016/S0968-0004(01)02027-8. URL https://linkinghub.elsevier.com/retrieve/pii/S0968000401020278.

36. Jaime Huerta-Cepas, Joaquín Dopazo, and Toni Gabaldón. ETE: a python Environment for Tree Exploration. BMC Bioinformatics, 11(1):24, December 2010. ISSN 1471-2105. doi: 10.1186/1471-2105-11-24. URL https://bmcbioinformatics.biomedcentral.com/articles/10.1186/1471-2105-11-24.

37. The UniProt Consortium, Alex Bateman, Maria-Jesus Martin, Sandra Orchard, Michele Magrane, Rahat Agivetova, Shadab Ahmad, Emanuele Alpi, Emily H Bowler-Barnett, Ramona Britto, Borisas Bursteinas, Hema Bye-A-Jee, Ray Coetzee, Austra Cukura, Alan Da Silva, Paul Denny, Tunca Dogan, ThankGod Ebenezer, Jun Fan, Leyla Garcia Castro, Penelope Garmiri, George Georghiou, Leonardo Gonzales, Emma Hatton-Ellis, Abdulrahman Hussein, Alexandr Ignatchenko, Giuseppe Insana, Rizwan Ishtiaq, Petteri Jokinen, Vishal Joshi, Dushyanth Jyothi, Antonia Lock, Rodrigo Lopez, Aurelien Luciani, Jie Luo, Yvonne Lussi, Alistair MacDougall, Fabio Madeira, Mahdi Mahmoudy, Manuela Menchi, Alok Mishra, Katie Moulang, Andrew Nightingale, Carla Susana Oliveira, Sangya Pundir, Guoying Qi, Shriya Raj, Daniel Rice, Milagros Rodriguez Lopez, Rabie Saidi, Joseph Sampson, Tony Sawford, Elena Speretta, Edward Turner, Nidhi Tyagi, Preethi Vasudev, Vladimir Volynkin, Kate Warner, Xavier Watkins, Rossana Zaru, Hermann Zellner, Alan Bridge, Sylvain Poux, Nicole Redaschi, Lucila Aimo, Ghislaine Argoud-Puy, Andrea Auchincloss, Kristian Axelsen, Parit Bansal, Delphine Baratin, Marie-Claude Blatter, Jerven Bolleman, Emmanuel Boutet, Lionel Breuza, Cristina Casals-Casas, Edouard De Castro, Kamal Chikh Echioukh, Elisabeth Coudert, Beatrice Cuche, Mikael Doche, Dolnide Dornevil, Anne Estreicher, Maria Livia Famiglietti, Marc Feuermann, Elisabeth Gasteiger, Sebastien Gehant, Vivienne Gerritsen, Arnaud Gos, Nadine Gruaz-Gumowski, Ursula Hinz, Chantal Hulo, Nevila Hyka-Nouspikel, Florence Jungo, Guillaume Keller, Arnaud Kerhornou, Vicente Lara, Philippe Le Mercier, Damien Lieberherr, Thierry Lombardot, Xavier Martin, Patrick Masson, Anne Morgat, Teresa Batista Neto, Salvo Paesano, Ivo Pedruzzi, Sandrine Pilbout, Lucille Pourcel, Monica Pozzato, Manuela Pruess, Catherine Rivoire, Christian Sigrist, Karin Sonesson, Andre Stutz, Shyamala Sundaram, Michael Tognolli, Laure Verbregue, Cathy H Wu, Cecilia N Arighi, Leslie Arminski, Chuming Chen, Yongxing Chen, John S Garavelli, Hongzhan Huang, Kati Laiho, Peter McGarvey, Darren A Natale, Karen Ross,R R Vinayaka, Qinghua Wang, Yuqi Wang, Lai-Su Yeh, Jian Zhang, Patrick Ruch, and Douglas Teodoro. UniProt: the universal protein knowledgebase in 2021. Nucleic Acids Research, 49 (D1):D480–D489, January 2021. ISSN 0305-1048, 1362-4962. doi: 10.1093/nar/gkaa1100. URL https://academic.oup.com/nar/article/49/D1/D480/6006196.

38. Martin Steinegger and Johannes Söding. MMseqs2 enables sensitive protein sequence searching for the analysis of massive data sets. Nature Biotechnology, 35(11):1026–1028, November 2017. ISSN 1087-0156, 1546-1696. doi: 10.1038/nbt.3988. URL https://www.nature.com/articles/nbt.3988.

39. Zeming Lin, Halil Akin, Roshan Rao, Brian Hie, Zhongkai Zhu, Wenting Lu, Nikita Smetanin, Robert Verkuil, Ori Kabeli, Yaniv Shmueli, Allan Dos Santos Costa, Maryam Fazel-Zarandi, Tom Sercu, Salvatore Candido, and Alexander Rives. Evolutionary-scale prediction of atomic-level protein structure with a language model. Science, 379(6637):1123–1130, March 2023. ISSN 0036-8075, 1095-9203. doi: 10.1126/science.ade2574. URL https://www.science.org/doi/10.1126/science.ade2574.

40. Alexander Rives, Joshua Meier, Tom Sercu, Siddharth Goyal, Zeming Lin, Jason Liu, Demi Guo, Myle Ott, C. Lawrence Zitnick, Jerry Ma, and Rob Fergus. Biological structure and function emerge from scaling unsupervised learning to 250 million protein sequences. Proceedings of the National Academy of Sciences, 118(15):e2016239118, April 2021. ISSN 0027-8424, 1091-6490. doi: 10.1073/pnas.2016239118. URL https://pnas.org/doi/full/10.1073/pnas.2016239118.

41. Ahmed Elnaggar, Michael Heinzinger, Christian Dallago, Ghalia Rehawi, Yu Wang, Llion Jones, Tom Gibbs, Tamas Feher, Christoph Angerer, Martin Steinegger, Debsindhu Bhowmik, and Burkhard Rost. ProtTrans: Toward Understanding the Language of Life Through Self-Supervised Learning. IEEE Transactions on Pattern Analysis and Machine Intelligence, 44 (10):7112–7127, October 2022. ISSN 0162-8828, 2160-9292, 1939-3539. doi: 10.1109/TPAMI.2021.3095381. URL https://ieeexplore.ieee.org/document/9477085/.

42. Ahmed Elnaggar, Hazem Essam, Wafaa Salah-Eldin, Walid Moustafa, Mohamed Elkerdawy, Charlotte Rochereau, and Burkhard Rost. Ankh: Optimized Protein Language Model Unlocks General-Purpose Modelling, 2023. URL https://arxiv.org/abs/2301.06568. Version Number: 1.

43. Michael Heinzinger, Ahmed Elnaggar, Yu Wang, Christian Dallago, Dmitrii Nechaev, Florian Matthes, and Burkhard Rost. Modeling aspects of the language of life through transfer-learning protein sequences. BMC Bioinformatics, 20(1):723, December 2019. ISSN 1471-2105. doi:10.1186/s12859-019-3220-8. URL https://bmcbioinformatics.biomedcentral.com/articles/10.1186/s12859-019-3220-8.

44. Ethan C. Alley, Grigory Khimulya, Surojit Biswas, Mohammed AlQuraishi, and George M. Church. Unified rational protein engineering with sequence-based deep representation learning. Nature Methods, 16(12):1315–1322, December 2019. ISSN 1548-7091, 1548-7105. doi: 10.1038/s41592-019-0598-1. URL https://www.nature.com/articles/s41592-019-0598-1.

45. Seonwoo Min, Seunghyun Park, Siwon Kim, Hyun-Soo Choi, Byunghan Lee, and Sungroh Yoon. Pre-Training of Deep Bidirectional Protein Sequence Representations with Structural Information, 2019. URL https://arxiv.org/abs/1912.05625. Version Number: 4.

46. Amy X. Lu, Haoran Zhang, Marzyeh Ghassemi, and Alan Moses. Self-Supervised Contrastive Learning of Protein Representations By Mutual Information Maximization, September 2020. URL http://biorxiv.org/lookup/doi/10.1101/2020.09.04.283929.

47. Piotr Bojanowski, Edouard Grave, Armand Joulin, and Tomas Mikolov. Enriching Word Vectors with Subword Information. Transactions of the Association for Computational Linguistics, 5: 135–146, December 2017. ISSN 2307-387X. doi: 10.1162/tacl_a_00051. URL https://direct.mit.edu/tacl/article/43387.

48. Jeffrey Pennington, Richard Socher, and Christopher Manning. Glove: Global Vectors for Word Representation. In Proceedings of the 2014 Conference on Empirical Methods in Natural Language Processing (EMNLP), pages 1532–1543, Doha, Qatar, 2014. Association for Computational Linguistics. doi: 10.3115/v1/D14-1162. URL http://aclweb.org/anthology/D14-1162.

49. Tomas Mikolov, Ilya Sutskever, Kai Chen, Greg S Corrado, and Jeff Dean. Distributed representations of words and phrases and their compositionality. In C.J. Burges, L. Bottou, M. Welling, Z. Ghahramani, and K.Q. Weinberger, editors, Advances in neural information processing systems, volume 26. Curran Associates, Inc., 2013. URL https://proceedings.neurips.cc/paper_files/paper/2013/file/9aa42b31882ec039965f3c4923ce901b-Paper.pdf.

50. Takuya Akiba, Shotaro Sano, Toshihiko Yanase, Takeru Ohta, and Masanori Koyama. Optuna: A Next-generation Hyperparameter Optimization Framework, 2019. URL https://arxiv.org/abs/1907.10902. Version Number: 1.

51. Philipp Koehn. Statistical significance tests for machine translation evaluation. In Proceedings of the 2004 conference on empirical methods in natural language processing, pages 388–395, 2004.

52. Taylor Berg-Kirkpatrick, David Burkett, and Dan Klein. An empirical investigation of statistical significance in NLP. In Proceedings of the 2012 joint conference on empirical methods in natural language processing and computational natural language learning, pages 995–1005, 2012.

53. Yoav Benjamini and Yosef Hochberg. Controlling the False Discovery Rate: A Practical and Powerful Approach to Multiple Testing. Journal of the Royal Statistical Society Series B: Statistical Methodology, 57(1):289–300, January 1995. ISSN 1369-7412, 1467-9868. doi: 10.1111/j.2517-6161.1995.tb02031.x. URL https://academic.oup.com/jrsssb/article/57/1/289/7035855.

