## Supplementary Tables and Figures for "AdventML: Advanced Enzyme Temperature Prediction with Transformer-Based Embeddings and Resampling Strategies"

### Supplementary Information

**Table S1. Effect of OGT features on prediction performance across feature sets.** Evaluated on the original TOMER train-test split (2,917 samples). Performance values are averaged over HistGradientBoosting, CatBoost, and SVR, each optimised with 96 Optuna trials. PLM embeddings were obtained via average pooling.

| | OGT | | | no OGT | | | $\lvert\Delta$ OGT - no OGT$\rvert$ | | |
|---|---|---|---|---|---|---|---|---|---|
| Protein Representations | MAE | RMSE | $R^2$ | MAE | RMSE | $R^2$ | MAE | RMSE | $R^2$ |
| esm3_sm_open_v1 | **8.61** | **11.53** | **0.605** | **9.44** | **12.55** | **0.533** | 0.83 | 1.02 | 0.073 |
| esmc_300m | 8.95 | 11.95 | 0.571 | **9.47** | **12.54** | **0.528** | **0.52** | **0.59** | **0.043** |
| **ProtTransT5XLU50** | 8.99 | **11.95** | 0.575 | 9.60 | 12.65 | **0.524** | 0.61 | 0.70 | 0.051 |
| esmc_600m | 9.48 | 12.92 | 0.547 | 9.87 | 13.39 | 0.513 | **0.39** | **0.47** | **0.033** |
| ProtTransT5UniRef50 | **8.79** | 11.99 | 0.571 | 9.53 | 12.89 | 0.505 | 0.74 | 0.90 | 0.066 |
| ProtTransT5BFD | 8.94 | 12.22 | 0.554 | 9.58 | 13.03 | 0.493 | 0.64 | 0.81 | 0.061 |
| ProtTransBertBFD | **8.50** | **11.20** | 0.566 | **9.49** | **12.24** | 0.483 | 0.99 | 1.03 | 0.084 |
| prime | 9.98 | 13.13 | 0.483 | 10.03 | 13.22 | 0.476 | **0.05** | **0.09** | **0.007** |
| ProtTransAlbertBFD | 9.41 | 12.63 | 0.563 | 10.62 | 14.08 | 0.457 | 1.21 | 1.45 | 0.106 |
| ProtTransXLNetUniRef100 | 9.31 | 12.59 | 0.556 | 10.55 | 14.02 | 0.450 | 1.24 | 1.43 | 0.106 |
| FastText | 9.40 | 12.50 | **0.583** | 11.57 | 15.25 | 0.380 | 2.17 | 2.75 | 0.202 |
| SeqVec | 9.05 | 12.18 | 0.558 | 11.12 | 14.55 | 0.372 | 2.08 | 2.38 | 0.186 |
| CPCProt | 10.34 | 13.29 | 0.550 | 12.40 | 15.85 | 0.363 | 2.06 | 2.57 | 0.187 |
| Rostlab_ProstT5 | 9.44 | 12.53 | 0.529 | 11.34 | 14.74 | 0.347 | 1.90 | 2.21 | 0.181 |
| Glove | 9.56 | 12.53 | 0.522 | 11.41 | 14.84 | 0.330 | 1.85 | 2.31 | 0.192 |
| Bepler | 9.26 | 12.11 | **0.580** | 11.76 | 15.32 | 0.329 | 2.50 | 3.20 | 0.251 |
| PLUSRNN | 10.49 | 13.91 | 0.514 | 12.67 | 16.69 | 0.304 | 2.18 | 2.78 | 0.210 |
| aaindex | 9.09 | 12.04 | 0.504 | 11.22 | 14.41 | 0.291 | 2.14 | 2.37 | 0.214 |
| composition | 10.83 | 14.20 | 0.436 | 12.59 | 16.36 | 0.257 | 1.76 | 2.16 | 0.179 |
| simple | 9.76 | 13.50 | 0.408 | 12.19 | 16.01 | 0.168 | 2.42 | 2.52 | 0.240 |
| physchem | 9.79 | 12.91 | 0.518 | 13.91 | 17.49 | 0.118 | 4.11 | 4.57 | 0.400 |
| auto | 10.20 | 13.60 | 0.420 | 14.69 | 18.77 | - | 4.49 | 5.17 | - |

**Table S5. Comparison of SOTA models on the held-out test set.**

| Model | $R^2$ | MAE | RMSE | MAE $>90$ | MAE 70-90 | MAE $<30$ |
|---|---|---|---|---|---|---|
| **AdventML** | **0.646** | **7.58** | **10.14** | 16.78 | 11.32 | **8.04** |
| SegmentTransformer (retrain) | **0.606** | **7.68** | **10.70** | 22.41 | 12.47 | 9.47 |
| TOMER & Pro-Prime | **0.605** | **8.28** | **10.72** | **8.58** | **8.13** | 13.75 |
| AdventML (tail) | 0.558 | 8.45 | 11.33 | **12.33** | **9.48** | **7.42** |
| AdventML (extreme) | 0.536 | 8.85 | 11.61 | **12.77** | **9.16** | **8.78** |
| Seq2Topt | 0.455 | 9.36 | 12.59 | 18.12 | 13.44 | 12.88 |
| Seq2Topt (retrain) | 0.443 | 9.57 | 12.73 | 27.05 | 20.56 | 9.18 |
| TOMER | 0.079 | 11.31 | 16.37 | 54.17 | 35.12 | 13.49 |
| SegmentTransformer | -0.190 | 15.29 | 18.60 | 13.11 | 12.16 | 23.16 |

**Table S2. Comparison of protein representations on the evaluation set.** Sequence-level PLM embeddings were obtained via average pooling. For each representation, seven models (HistGradientBoosting, XGBoost, CatBoost, QuantileGBR, ElasticNet, SVR, and Kernel-Ridge) were trained with hyperparameters optimised over 96 Optuna trials. Reported values are averaged across all models.

| Protein Representation | $R^2$ | MAE | MAE >90 | MAE 70-90 | MAE <30 |
|---|---|---|---|---|---|
| **PT T5 XL U50** | **0.530** | **7.93** | **21.80** | 15.32 | **9.77** |
| Ankh3 XL | **0.514** | 8.14 | **20.54** | 17.14 | 10.49 |
| ESMC 600M | **0.512** | **8.10** | 22.86 | **13.64** | 10.22 |
| PT T5 U50 | 0.502 | **7.98** | 23.06 | 15.26 | 9.99 |
| ESM2 650M | 0.494 | 8.19 | **19.67** | 17.00 | 10.02 |
| ESM2 3B | 0.490 | 8.25 | 23.42 | 17.02 | **9.73** |
| ESMC 300M | 0.486 | 8.22 | 23.42 | **14.88** | **9.77** |
| Prime | 0.479 | 8.27 | 26.05 | **13.09** | 11.93 |
| PT T5 BFD | 0.453 | 8.22 | 28.70 | 16.07 | 10.51 |
| ESM2 35M | 0.450 | 13.19 | 26.57 | 23.68 | 13.81 |
| Ankh Large | 0.441 | 8.73 | 29.98 | 18.66 | 9.91 |
| PT Bert BFD | 0.431 | 8.60 | 28.65 | 16.89 | 11.25 |
| Ankh3 Large | 0.429 | 8.69 | 29.32 | 19.64 | 11.51 |
| PT Albert BFD | 0.420 | 8.31 | 26.35 | 18.95 | 10.65 |
| PT XLNet U100 | 0.405 | 8.84 | 28.79 | 20.83 | 10.64 |
| Ankh Base | 0.400 | 13.40 | 35.19 | 22.09 | 13.53 |
| PLUS RNN | 0.368 | 9.21 | 29.12 | 20.18 | 11.12 |
| SeqVec | 0.345 | 9.51 | 33.11 | 20.86 | 12.27 |
| *Simple | 0.312 | 14.49 | 38.01 | 27.68 | 14.65 |
| *AAIndex | 0.291 | 14.45 | 41.10 | 29.62 | 14.65 |
| GloVe | 0.288 | 13.99 | 41.13 | 28.54 | 14.63 |
| FastText | 0.286 | 14.42 | 42.64 | 28.31 | 15.31 |
| CPCProt | 0.277 | 14.49 | 41.98 | 28.59 | 15.33 |
| *Composition | 0.276 | 14.26 | 41.62 | 27.53 | 15.00 |
| Bepler | 0.250 | 14.13 | 37.66 | 29.93 | 14.40 |
| *Physicochemical | 0.160 | 15.17 | 51.70 | 31.95 | 16.48 |
| *Auto | 0.033 | 16.33 | 56.57 | 38.95 | 16.40 |

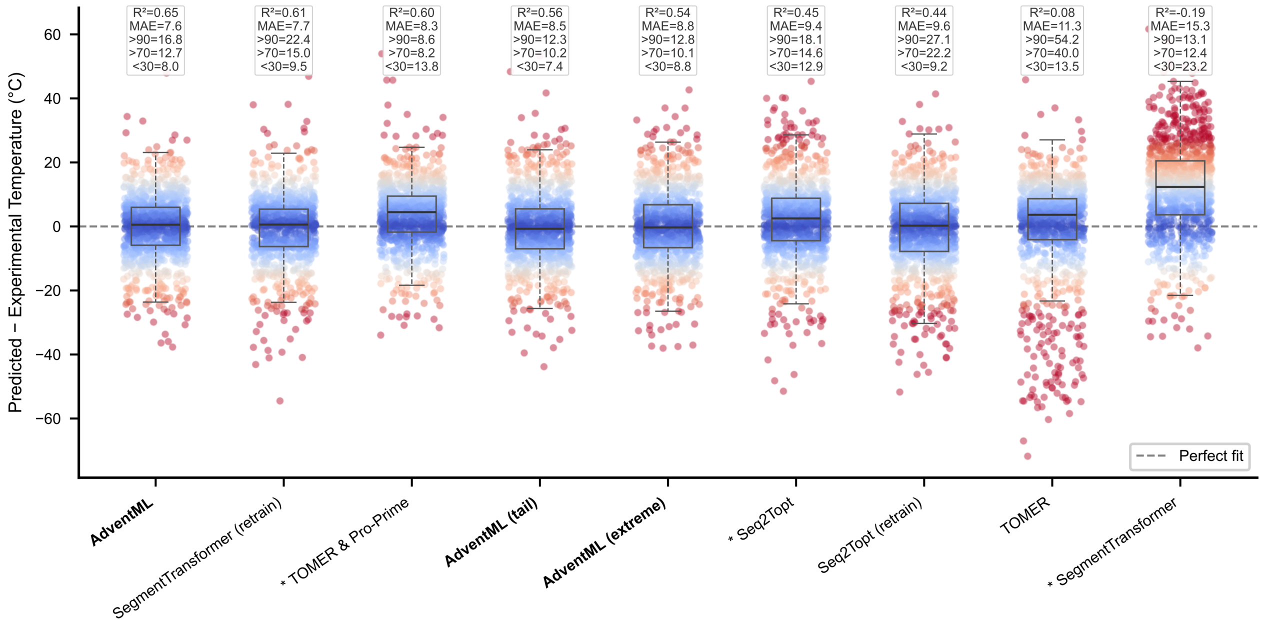


**Figure S1. Residual distribution by temperature bin.** Strip plot of prediction residuals for AdventML and all benchmarked models on the held-out test set. Positive values correspond to predictions exceeding the experimental $T_{opt}$. Asterisks (*) denote models whose training data overlaps with the test set.

**Table S3.** Comparison of resampling strategies.

| Resampling Method | $R^2$ | MAE | $\text{MAE}_{\text{tail}}$ | MAE $>60$ | MAE $<25$ |
|---|---|---|---|---|---|
| Tomek Links | **0.456** | **8.43** | 14.96 | 15.43 | 14.21 |
| ADASYN | **0.457** | **8.48** | 14.75 | 15.16 | 14.08 |
| EXT TomekLinks | 0.446 | **8.50** | 14.33 | 14.36 | 14.28 |
| **Baseline** | **0.447** | 8.53 | 14.70 | 14.82 | 14.50 |
| EXT KNNOR | 0.430 | 8.66 | 13.91 | 13.63 | 14.33 |
| KNNOR | 0.428 | 8.67 | 14.19 | 13.97 | 14.53 |
| EXT SMOGN | 0.414 | 8.78 | 13.37 | 13.08 | **13.80** |
| EXT SMOTER | 0.412 | 8.81 | 13.40 | 13.13 | 13.81 |
| EXT ADASYN | 0.417 | 8.83 | 13.39 | 13.40 | **13.35** |
| EXT WERCS | 0.398 | 8.88 | 13.43 | 12.87 | 14.30 |
| EXT WERCS-GN | 0.397 | 8.91 | 13.39 | 12.77 | 14.33 |
| IHT | 0.396 | 8.93 | 19.38 | 21.54 | 16.00 |
| WERCS-GN | 0.399 | 8.99 | 13.53 | 13.02 | 14.31 |
| WERCS | 0.397 | 9.00 | 13.63 | 13.20 | 14.30 |
| Cluster Centroids | 0.405 | 9.02 | 16.00 | 16.40 | 15.38 |
| SMOGN | 0.383 | 9.04 | 13.53 | 12.78 | 14.70 |
| SMOTER | 0.382 | 9.05 | 13.62 | 12.87 | 14.78 |
| **EXT ENN** | 0.358 | 9.19 | **12.90** | **12.29** | 13.85 |
| CNN | 0.379 | 9.21 | 15.97 | 16.11 | 15.75 |
| EXT Random Over | 0.353 | 9.22 | 13.26 | 12.65 | 14.20 |
| EXT Cluster Centroids | 0.365 | 9.24 | 13.28 | 12.76 | 14.07 |
| EXT IHT | 0.341 | 9.25 | 13.18 | **12.13** | 14.80 |
| EXT CNN | 0.354 | 9.29 | 13.10 | 12.48 | 14.04 |
| EXT Random Under | 0.351 | 9.29 | **13.05** | **12.36** | 14.12 |
| ENN | 0.280 | 9.66 | 21.90 | 25.62 | 16.09 |
| EXT OSS | 0.317 | 9.82 | 14.46 | 14.41 | 14.52 |
| EXT NearMiss | 0.243 | 10.23 | **13.09** | 12.50 | 14.00 |
| NearMiss | 0.259 | 10.35 | 17.80 | 17.39 | 18.42 |
| OSS | 0.235 | 10.48 | 13.12 | 14.01 | **11.71** |

**Table S4. Comparison of all trained models on the eval set.**

| | Training | | Eval | | $\|\Delta$ Train – Eval$\|$ | | Eval MAE (ranges) | | |
|---|---|---|---|---|---|---|---|---|---|
| Model | $R^2$ | MAE | $R^2$ | MAE | $\Delta R^2$ | ΔMAE | $> 90$ | $< 30$ | tail |
| GPR | **1.000** | **0.00** | **0.560** | **7.66** | 0.440 | 7.66 | 21.41 | 9.11 | 16.27 |
| KernelRidge | **0.941** | **2.41** | **0.559** | **7.50** | 0.382 | 5.09 | 17.17 | 8.38 | 15.80 |
| BaggingMLP | 0.832 | 4.55 | **0.534** | **7.82** | 0.298 | 3.27 | 17.24 | 7.17 | 12.50 |
| ResNetMLP | **0.915** | **3.34** | 0.524 | 7.84 | 0.391 | 4.50 | 14.99 | 6.59 | 14.42 |
| MLP | 0.708 | 6.12 | 0.510 | 8.22 | 0.198 | 2.10 | 16.68 | 9.53 | 13.22 |
| RandomForest | 0.764 | 5.56 | 0.489 | 8.43 | 0.276 | 2.87 | 20.40 | 8.56 | 14.82 |
| ARDRegression | 0.582 | 7.47 | 0.489 | 8.37 | 0.094 | 0.90 | 23.48 | 8.95 | 16.89 |
| RealMLP | 0.758 | 5.48 | 0.486 | 8.24 | 0.272 | 2.76 | 17.20 | 6.42 | 12.78 |
| ExtraTrees | 0.742 | 5.78 | 0.471 | 8.61 | 0.271 | 2.83 | 19.15 | 8.04 | 13.94 |
| WideDeep | 0.847 | 4.53 | 0.471 | 8.46 | 0.376 | 3.93 | 15.79 | 6.92 | 13.20 |
| SparseMLP | 0.830 | 4.71 | 0.465 | 8.46 | 0.365 | 3.75 | 16.23 | 7.51 | 13.35 |
| PLS | 0.595 | 7.40 | 0.458 | 8.63 | 0.137 | 1.23 | 22.87 | 8.75 | 16.49 |
| TabM | 0.795 | 4.63 | 0.453 | 8.39 | 0.342 | 3.75 | 17.73 | 7.56 | 12.70 |
| OMP | 0.491 | 8.28 | 0.452 | 8.74 | 0.039 | 0.46 | 25.91 | 9.74 | 17.99 |
| **CatBoost** | 0.708 | 5.97 | 0.449 | 8.84 | 0.259 | 2.86 | **13.28** | 7.42 | 12.39 |
| HistGradientBoosting | 0.664 | 6.57 | 0.446 | 8.90 | 0.218 | 2.33 | 16.07 | 7.59 | 12.69 |
| DANets | 0.548 | 7.99 | 0.444 | 8.92 | 0.104 | 0.92 | 15.99 | 6.87 | 12.95 |
| PassiveAggressive | 0.572 | 7.17 | 0.442 | 8.56 | 0.130 | 1.39 | 21.58 | 8.28 | 16.21 |
| BaselineMLP | 0.738 | 6.03 | 0.438 | 8.64 | 0.300 | 2.61 | 13.52 | 6.75 | 12.49 |
| Node | 0.558 | 7.82 | 0.437 | 8.93 | 0.121 | 1.11 | 18.08 | **5.66** | 12.75 |
| **BaggingSVR** | 0.651 | 6.39 | 0.436 | 9.05 | 0.215 | 2.65 | **13.19** | 7.22 | 12.12 |
| GANDALF | 0.573 | 7.55 | 0.436 | 8.85 | 0.137 | 1.30 | 19.09 | 7.65 | 12.62 |
| LassoNet | 0.594 | 7.41 | 0.436 | 8.85 | 0.159 | 1.44 | 14.42 | 7.73 | 11.98 |
| TabR | 0.681 | 6.42 | 0.435 | 8.68 | 0.246 | 2.26 | 15.74 | 6.18 | 12.10 |
| BaggingDecisionTree | 0.629 | 7.21 | 0.429 | 9.06 | 0.200 | 1.86 | 18.10 | 8.00 | 13.75 |
| XGBoost | 0.654 | 6.80 | 0.420 | 9.14 | 0.234 | 2.34 | 13.86 | 7.11 | 12.23 |
| GradientBoosting | 0.694 | 5.92 | 0.419 | 9.08 | 0.274 | 3.16 | 13.60 | 6.80 | 12.30 |
| AdaBoost | 0.460 | 8.77 | 0.419 | 9.09 | 0.041 | **0.32** | 21.29 | 12.03 | 15.07 |
| SVR | 0.634 | 6.00 | 0.418 | 9.13 | 0.216 | 3.13 | 13.58 | 6.92 | 12.54 |
| FTTransformer | 0.432 | 8.60 | 0.417 | 8.78 | **0.015** | **0.18** | 19.87 | 8.44 | 14.13 |
| GatedMLP | 0.748 | 5.93 | 0.416 | 8.99 | 0.332 | 3.06 | 14.95 | 7.20 | 13.15 |
| RANSAC | 0.586 | 6.96 | 0.409 | 8.90 | 0.177 | 1.94 | 21.05 | 7.98 | 16.49 |
| TabNetInspired | 0.797 | 5.10 | 0.408 | 9.04 | 0.388 | 3.93 | 21.37 | 8.05 | 17.20 |
| AttentionMLP | 0.738 | 6.19 | 0.402 | 9.37 | 0.336 | 3.18 | 15.81 | 6.38 | 13.86 |
| Hopular | 0.565 | 7.60 | 0.399 | 9.08 | 0.165 | 1.48 | 15.70 | 6.67 | 11.95 |
| BaggingSVRExtreme | 0.593 | 6.92 | 0.386 | 9.51 | 0.208 | 2.59 | 13.51 | 7.37 | 12.16 |
| BaggingDecisionTreeExtreme | 0.584 | 7.51 | 0.375 | 9.56 | 0.209 | 2.05 | 16.71 | 7.60 | 12.97 |
| BaggingMLPExtreme | 0.626 | 6.85 | 0.366 | 9.54 | 0.260 | 2.69 | 13.69 | **5.59** | **11.23** |
| RTDLResNet | 0.480 | 8.24 | 0.356 | 9.31 | 0.124 | 1.06 | 14.94 | **5.36** | **11.18** |
| TabNet | 0.440 | 8.68 | 0.351 | 9.35 | 0.088 | 0.67 | 22.51 | 6.18 | 15.47 |
| BayesianRidge | 0.387 | 9.34 | 0.349 | 9.74 | **0.038** | 0.40 | 15.55 | 7.65 | 12.07 |
| SAINT | 0.367 | 9.29 | 0.342 | 9.55 | **0.025** | **0.26** | 27.38 | 6.74 | 17.81 |
| Ridge | 0.386 | 9.36 | 0.318 | 10.03 | 0.068 | 0.67 | 15.20 | 7.45 | 11.99 |
| LinearRegression | 0.258 | 10.33 | 0.093 | 11.56 | 0.165 | 1.23 | 15.61 | 9.35 | 13.03 |
| ElasticNet | 0.324 | 9.74 | 0.066 | 11.94 | 0.259 | 2.20 | 14.81 | 9.37 | 13.91 |
| Quantile | 0.158 | 10.29 | 0.064 | 11.58 | 0.094 | 1.29 | **10.74** | 10.50 | **10.52** |
| DecisionTree | 0.148 | 10.81 | -0.060 | 11.92 | 0.208 | 1.11 | 21.08 | 11.18 | 16.22 |
| Huber | 0.163 | 10.52 | -0.195 | 13.47 | 0.359 | 2.95 | 16.67 | 10.86 | 15.03 |
| SNN | -0.172 | 12.96 | -0.518 | 14.72 | 0.346 | 1.76 | 19.93 | 14.98 | 24.45 |

**Table S6. Pairwise MAE $>$70 significance matrix** (paired bootstrap, $B = 100{,}000$, BH-corrected $\alpha = 0.05$). $>$ row significantly better than column; $<$ significantly worse; = not significant. Models ordered by MAE $>$70 (ascending). Only p-values higher than 0.001 are shown.

| Model | ***TOMER & Pro-Prime*** | ***AdventML (extreme)*** | ***AdventML (tail)*** | ***SegmentTransformer*** | ***AdventML*** | ***Seq2Topt*** | ***SegmentTransformer (retrain)*** | ***Seq2Topt (retrain)*** | ***TOMER*** |
|---|---|---|---|---|---|---|---|---|---|
| **TOMER & Pro-Prime** | - | $>$ | $>$ | $>$ | $>$ | $>$ | $>$ | $>$ | $>$ |
| **AdventML (extreme)** | $<$ | - | = (0.70) | = (0.21) | $>$ | $>$ | $>$ | $>$ | $>$ |
| **AdventML (tail)** | $<$ | = (0.70) | - | = (0.26) | $>$ | $>$ | $>$ | $>$ | $>$ |
| **SegmentTransformer** | $<$ | = (0.21) | = (0.26) | - | = (0.23) | $>$ (0.01) | $>$ | $>$ | $>$ |
| **AdventML** | $<$ | $<$ | $<$ | = (0.23) | - | $>$ (0.04) | $>$ | $>$ | $>$ |
| **Seq2Topt** | $<$ | $<$ | $<$ | $<$ (0.01) | $<$ (0.04) | - | = (0.68) | $>$ | $>$ |
| **SegmentTransformer (retrain)** | $<$ | $<$ | $<$ | $<$ | $<$ | = (0.68) | - | $>$ | $>$ |
| **Seq2Topt (retrain)** | $<$ | $<$ | $<$ | $<$ | $<$ | $<$ | $<$ | - | $>$ |
| **TOMER** | $<$ | $<$ | $<$ | $<$ | $<$ | $<$ | $<$ | $<$ | - |

**Table S7. Pairwise MAE $>$90 significance matrix** (paired bootstrap, $B = 100{,}000$, BH-corrected $\alpha = 0.05$). $>$ row significantly better than column; $<$ significantly worse; = not significant. Models ordered by MAE $>$90 (ascending). Only p-values higher than 0.001 are shown.

| Model | ***TOMER & Pro-Prime*** | ***AdventML (tail)*** | ***AdventML (extreme)*** | ***SegmentTransformer*** | ***AdventML*** | ***Seq2Topt*** | ***SegmentTransformer (retrain)*** | ***Seq2Topt (retrain)*** | ***TOMER*** |
|---|---|---|---|---|---|---|---|---|---|
| **TOMER & Pro-Prime** | - | = (0.27) | = (0.07) | = (0.06) | $>$ | $>$ | $>$ | $>$ | $>$ |
| **AdventML (tail)** | = (0.27) | - | = (0.15) | = (0.50) | $>$ | = (0.13) | $>$ | $>$ | $>$ |
| **AdventML (extreme)** | = (0.07) | = (0.15) | - | = (0.81) | = (0.19) | = (0.25) | $>$ | $>$ | $>$ |
| **SegmentTransformer** | = (0.06) | = (0.50) | = (0.81) | - | = (0.61) | = (0.52) | $>$ (0.03) | $>$ | $>$ |
| **AdventML** | $<$ | $<$ | = (0.19) | = (0.61) | - | = (0.66) | $>$ | $>$ | $>$ |
| **Seq2Topt** | $<$ | = (0.13) | = (0.25) | = (0.52) | = (0.66) | - | = (0.27) | = (0.05) | $>$ |
| **SegmentTransformer (retrain)** | $<$ | $<$ | $<$ | $<$ (0.03) | $<$ | = (0.27) | - | = (0.15) | $>$ |
| **Seq2Topt (retrain)** | $<$ | $<$ | $<$ | $<$ | $<$ | = (0.05) | = (0.15) | - | $>$ |
| **TOMER** | $<$ | $<$ | $<$ | $<$ | $<$ | $<$ | $<$ | $<$ | - |

**Table S8. Pairwise MAE <30 significance matrix** (paired bootstrap, $B = 100{,}000$, BH-corrected $\alpha = 0.05$). $>$ row significantly better than column; $<$ significantly worse; = not significant. Models ordered by MAE <30 (ascending). Only p-values higher than 0.001 are shown.

| Model | AdventML (tail) | AdventML | AdventML (extreme) | Seq2Topt (retrain) | SegmentTransformer (retrain) | Seq2Topt | TOMER | TOMER & Pro-Prime | SegmentTransformer |
|---|---|---|---|---|---|---|---|---|---|
| **AdventML (tail)** | - | $>$ (0.05) | $>$ | $>$ | $>$ | $>$ | $>$ | $>$ | $>$ |
| **AdventML** | $<$ (0.05) | - | = (0.06) | $>$ (0.03) | $>$ | $>$ | $>$ | $>$ | $>$ |
| **AdventML (extreme)** | $<$ | = (0.06) | - | = (0.55) | = (0.18) | $>$ | $>$ | $>$ | $>$ |
| **Seq2Topt (retrain)** | $<$ | $<$ (0.03) | = (0.55) | - | = (0.55) | $>$ | $>$ | $>$ | $>$ |
| **SegmentTransformer (retrain)** | $<$ | $<$ | = (0.18) | = (0.55) | - | $>$ | $>$ | $>$ | $>$ |
| **Seq2Topt** | $<$ | $<$ | $<$ | $<$ | $<$ | - | = (0.32) | = (0.16) | $>$ |
| **TOMER** | $<$ | $<$ | $<$ | $<$ | $<$ | = (0.32) | - | = (0.13) | $>$ |
| **TOMER & Pro-Prime** | $<$ | $<$ | $<$ | $<$ | $<$ | = (0.16) | = (0.13) | - | $>$ |
| **SegmentTransformer** | $<$ | $<$ | $<$ | $<$ | $<$ | $<$ | $<$ | $<$ | - |

**Table S9. Pairwise $R^2$ significance matrix** (paired bootstrap, $B = 100{,}000$, BH-corrected $\alpha = 0.05$). $>$ row significantly better than column; $<$ significantly worse; = not significant. Models ordered by $R^2$ (descending). Only p-values higher than 0.001 are shown.

| Model | AdventML | SegmentTransformer (retrain) | TOMER & Pro-Prime | AdventML (tail) | AdventML (extreme) | Seq2Topt | Seq2Topt (retrain) | TOMER | SegmentTransformer |
|---|---|---|---|---|---|---|---|---|---|
| **AdventML** | - | $>$ (0.02) | $>$ (0.03) | $>$ | $>$ | $>$ | $>$ | $>$ | $>$ |
| **SegmentTransformer (retrain)** | $<$ (0.02) | - | = (0.95) | $>$ (0.04) | $>$ | $>$ | $>$ | $>$ | $>$ |
| **TOMER & Pro-Prime** | $<$ (0.03) | = (0.95) | - | $>$ (0.04) | $>$ | $>$ | $>$ | $>$ | $>$ |
| **AdventML (tail)** | $<$ | $<$ (0.04) | $<$ (0.04) | - | = (0.19) | $>$ | $>$ | $>$ | $>$ |
| **AdventML (extreme)** | $<$ | $<$ | $<$ | = (0.19) | - | $>$ (0.02) | $>$ | $>$ | $>$ |
| **Seq2Topt** | $<$ | $<$ | $<$ | $<$ | $<$ (0.02) | - | = (0.74) | $>$ | $>$ |
| **Seq2Topt (retrain)** | $<$ | $<$ | $<$ | $<$ | $<$ | = (0.74) | - | $>$ | $>$ |
| **TOMER** | $<$ | $<$ | $<$ | $<$ | $<$ | $<$ | $<$ | - | $>$ |
| **SegmentTransformer** | $<$ | $<$ | $<$ | $<$ | $<$ | $<$ | $<$ | $<$ | - |

**Table S11.** Overview of resampling methods evaluated for regression imbalance correction

| Short Name | Full Name | Description |
|---|---|---|
| Baseline | Baseline | No resampling applied; data is used as-is with standard scaling. |

*Continued on next page*

| Short Name | Full Name | Description |
|---|---|---|
| ADASYN | Adaptive Synthetic Sampling | Generates synthetic minority samples adaptively, placing more near decision boundaries. |
| Cluster Centroids | Cluster Centroids | Undersamples the majority by replacing clusters of majority samples with their centroids. |
| CNN | Condensed Nearest Neighbours | Iteratively selects a subset of the majority class that correctly classifies all remaining samples via 1-NN. |
| ENN | Edited Nearest Neighbours | Removes majority samples whose class label disagrees with the majority vote of their $k$ nearest neighbours. |
| IHT | Instance Hardness Threshold | Removes instances that are hardest to classify correctly under cross-validated probability estimates. |
| KNNOR | K-Nearest Neighbour Oversampling for Regression | Generates synthetic samples by interpolating between minority instances and their $k$ nearest neighbours. |
| NearMiss | NearMiss | Selects majority samples closest to minority samples using one of three distance-based heuristics. |
| OSS | One-Sided Selection | Combines Tomek link removal with condensed nearest neighbours to clean the majority class boundary. |
| SMOGN | Synthetic Minority Oversampling with Gaussian Noise | Oversamples rare-target regions using SMOTE-style interpolation combined with Gaussian noise injection. |
| SMOTER | Synthetic Minority Oversampling for Regression | Applies SMOTE-style interpolation to generate synthetic samples in underrepresented target regions. |
| Tomek Links | Tomek Links | Removes majority-class members of Tomek link pairs lying on or near the decision boundary. |
| WERCS | Weighted Relevance-based Combination Strategy | Jointly over- and undersamples using relevance-based weighting of training instances. |
| WERCS-GN | WERCS with Gaussian Noise | Extends WERCS by adding Gaussian noise to synthetically generated oversampled instances. |
| *Extreme (Tail-Focused) Variants* | | |
| EXT ADASYN | Extreme ADASYN | ADASYN variant targeting only the extreme (tail) temperature regions of the target distribution. |
| EXT Cluster Centroids | Extreme Cluster Centroids | Cluster centroid undersampling applied selectively to the middle temperature range. |
| EXT CNN | Extreme CNN | Condensed nearest neighbours restricted to undersampling the non-tail temperature region. |
| EXT ENN | Extreme ENN | Edited nearest neighbours that cleans the middle temperature range while preserving tail instances. |
| EXT IHT | Extreme IHT | Instance hardness thresholding focused on removing hard-to-predict instances from the middle range. |
| EXT KNNOR | Extreme KNNOR | KNN-based oversampling restricted to generating synthetic samples in the tail temperature regions. |
| EXT NearMiss | Extreme NearMiss | NearMiss undersampling applied to the non-tail region to increase relative tail representation. |
| EXT OSS | Extreme OSS | One-sided selection targeting majority samples in the middle temperature range for removal. |
| EXT Random Over | Extreme Random Oversampling | Random duplication of training instances drawn from the tail temperature regions. |
| EXT Random Under | Extreme Random Undersampling | Random removal of training instances from the middle temperature range. |
| EXT SMOGN | Extreme SMOGN | SMOGN oversampling restricted to synthesising new samples in the extreme temperature tails. |
| EXT SMOTER | Extreme SMOTER | SMOTER interpolation applied exclusively within the tail temperature regions. |
| EXT TomekLinks | Extreme TomekLinks | Tomek link cleaning applied to the boundary between tail and non-tail temperature regions. |

*Continued on next page*

| Short Name | Full Name | Description |
|---|---|---|
| EXT WERCS | Extreme WERCS | Relevance-weighted resampling that boosts tail-region instances while down-weighting the middle range. |
| EXT WERCS-GN | Extreme WERCS-GN | Extreme WERCS with additional Gaussian noise on oversampled tail instances for diversity. |

**Table S12.** Hyperparameter search space for resampling strategies

| Parameter | Description | Tested Values |
|---|---|---|
| sampling_strategy | Target ratio of minority to majority samples after resampling. | $\{0.1, 0.2, 0.3, 0.5, 0.7, 1.0, 2.0, 3.0, 5.0\}$ |
| k / k_neighbors | Number of nearest neighbours for synthetic sample generation (ADASYN). | $\{3, 5\}$ |
| n_neighbors | Number of nearest neighbours for cleaning or oversampling. | $\{1, 3, 5, 7\}$ |
| tolerance | Distance tolerance for neighbour-based instance removal (CNN, ENN). | $\{0.0, 0.05, 0.1, 3.0, 5.0, 10.0\}$ |
| noise_std | Standard deviation of Gaussian noise added to synthetic samples (SMOGN, WERCS-GN). | $\{0.01, 0.02, 0.05, 0.1\}$ |
| voting | Strategy for computing cluster centroids (Cluster Centroids). | {soft, hard} |
| threshold_percentile | Percentile threshold for identifying boundary pairs (TomekLinks). | $\{30, 50, 70, 80, 90, 95\}$ |
| cv | Number of cross-validation folds for Instance Hardness Threshold. | $\{3, 5\}$ |
| rel_thres | Relevance function threshold for determining rare instances (ADASYN). | $\{0.2, 0.25\}$ |
| rel_coef | Relevance function coefficient controlling shape of the relevance curve. | $\{0.5, 1.0, 2.0, 5.0\}$ |
| samp_method | Sampling method for ADASYN (balance vs. extreme mode). | {balance, extreme} |
| n_seeds_S | Number of seed samples for condensed nearest neighbours (CNN, OSS). | $\{1, 3, 5\}$ |
| version | NearMiss algorithm version controlling the distance heuristic. | $\{1, 2, 3\}$ |
| bins | Number of bins for the relevance function discretisation (SMOGN, SMOTER). | $\{20, 40\}$ |
| critical_percentile | Percentile for identifying the critical zone boundary (OSS). | $\{0.05, 0.1\}$ |
| num_nbrs | Number of neighbours for KNN-based oversampling (KNNOR). | $\{3, 5\}$ |
| proportion_of_minority | Target proportion of minority instances after oversampling (KNNOR). | $\{0.7, 0.9\}$ |
| alpha | Interpolation coefficient for synthetic sample placement (KNNOR). | $\{0.25, 0.5, 0.7\}$ |

**Table S13. Deep learning architectures naming.**

| Model | Description |
|---|---|
| Attention MLP | Multi-head self-attention over features combined with a feed-forward MLP. |
| Baseline MLP | Standard multi-layer perceptron with configurable depth, dropout, and batch normalisation. |

*Continued on next page*

| Model | Description |
|---|---|
| DANets | Deep Abstract Networks that learn to group correlated input features into abstract layers. |
| FT-Transformer | Feature Tokenizer Transformer that converts each feature into a token processed by a Transformer encoder. |
| GANDALF | Gated Additive Neural Decision Forest that applies learned feature gates before soft decision trees. |
| Gated MLP | MLP preceded by per-feature sigmoid gates for soft feature selection. |
| GrowNet | Gradient boosting framework that uses shallow neural networks as weak learners in an additive ensemble. |
| Hopular | Modern Hopfield network layers that use the training set as associative memory for tabular lookup. |
| LassoNet | MLP with a linear skip connection and a hierarchy constraint that induces feature-level sparsity. |
| NODE | Neural Oblivious Decision Ensembles with differentiable oblivious decision trees and entmax splits. |
| RealMLP | Modern default-tuned MLP with LayerNorm, cosine-annealing schedule, and carefully chosen initialisation. |
| ResNet MLP | Dense layers with residual (skip) connections enabling deeper MLP training. |
| RTDL ResNet | Pre-activation residual MLP blocks following the Revisiting Deep Learning for Tabular Data design. |
| SAINT | Self-Attention and Intersample Attention Transformer operating on both features and samples. |
| SNN | Self-Normalising Neural Network using SELU activations and AlphaDropout for automatic normalisation. |
| Sparse MLP | MLP with group-sparse $\ell_{2,1}$ regularisation on the first layer for embedded feature selection. |
| TabM | Parameter-efficient ensembling of multiple tabular prediction heads within a single backbone. |
| TabNet | Sequential attentive feature selection with learnable sparsemax masks at each decision step. |
| TabNet-Inspired | Simplified attention-based sequential feature selector inspired by the TabNet architecture. |
| TabR | MLP encoder augmented with attention-based retrieval over nearest training neighbours. |
| VIME | Self- and semi-supervised pretraining via feature-mask estimation for tabular representation learning. |
| Wide & Deep | Combines a wide (linear) component for memorisation with a deep MLP for generalisation. |

**Table S14. Hyperparameter search spaces for predictive modeling pipelines**

| Pipeline | Hyperparameter | Search Space |
|---|---|---|
| *Linear Models* | | |
| **LinearRegression** | fit_intercept | {True, False} |
| | positive | {True, False} |
| **Ridge** | alpha | $10^{-6}, 10^{3}$ (log) |
| | solver | {auto, svd, cholesky, lsqr, sparse_cg, sag, saga} |
| | fit_intercept | {True, False} |
| **ElasticNet** | alpha | $10^{-5}, 10^{-2}$ (log) |
| | l1_ratio | $0.0, 1.0$ |
| | fit_intercept | {True, False} |
| | selection | {cyclic, random} |
| **BayesianRidge** | alpha_1 | $10^{-7}, 10^{-5}$ (log) |

*Continued on next page*

| Pipeline | Hyperparameter | Search Space |
|---|---|---|
| | alpha_2 | $10^{-7}, 10^{-5}$ (log) |
| | lambda_1 | $10^{-7}, 10^{-5}$ (log) |
| | lambda_2 | $10^{-7}, 10^{-5}$ (log) |
| | fit_intercept | {True, False} |
| **ARDRegression** | alpha_1 | $10^{-7}, 10^{-4}$ (log) |
| | alpha_2 | $10^{-7}, 10^{-4}$ (log) |
| | lambda_1 | $10^{-7}, 10^{-4}$ (log) |
| | lambda_2 | $10^{-7}, 10^{-4}$ (log) |
| | fit_intercept | {True, False} |
| **Huber** | epsilon | $1.01, 2.0$ |
| | alpha | $10^{-5}, 10^{-1}$ (log) |
| | fit_intercept | {True, False} |
| **PassiveAggressive** | C | $0.001, 10.0$ (log) |
| | epsilon | $0.01, 1.0$ |
| | loss | {epsilon_insensitive, squared_epsilon_insensitive} |
| | fit_intercept | {True, False} |
| **OMP** | n_nonzero_coefs | $1, 50$ |
| | fit_intercept | {True, False} |
| **PLS** | n_components | $1, 50$ |
| | scale | {True, False} |
| **Quantile** | quantile | $\{0.25, 0.5, 0.75\}$ |
| | alpha | $0.0, 1.0$ |
| | solver | {highs-ds, highs-ipm, highs} |
| | fit_intercept | {True, False} |
| **RANSAC** | min_samples | $0.1, 0.9$ |
| | residual_threshold | $1.0, 10.0$ |
| *Kernel Methods* | | |
| **SVR** | C | $0.01, 1000$ (log) |
| | epsilon | $0.001, 1.0$ (log) |
| | kernel | {linear, poly, rbf, sigmoid} |
| | gamma | {scale, auto} |
| | degree | $2, 5$ |
| **KernelRidge** | alpha | $0.01, 100$ (log) |
| | kernel | {linear, poly, rbf, sigmoid, laplacian} |
| | gamma | $0.001, 10.0$ (log) |
| | degree | $2, 5$ |
| | coef0 | $0.0, 1.0$ |
| **GPR** | kernel | {RBF, Matern, RationalQuadratic, DotProduct} |
| | length_scale | $10^{-3}, 100$ (log) |
| | alpha | $10^{-8}, 10^{-2}$ (log) |
| | n_restarts_optimizer | $0, 10$ |
| | normalize_y | {True, False} |
| *Scikit-learn Neural Network* | | |
| **MLP** | hidden_layer_sizes | {(50,), (100,), (50,50), (100,50), (100,100), (100,50,25)} |
| | activation | {relu, tanh, logistic} |
| | solver | {adam, sgd, lbfgs} |
| | alpha | $10^{-5}, 10^{-1}$ (log) |
| | learning_rate | {constant, invscaling, adaptive} |
| | learning_rate_init | $10^{-4}, 5 \times 10^{-3}$ (log) |
| *Decision Trees and Ensembles* | | |
| **DecisionTree** | max_depth | $2, 30$ |

*Continued on next page*

| Pipeline | Hyperparameter | Search Space |
|---|---|---|
| | min_samples_split | $2, 20$ |
| | min_samples_leaf | $1, 10$ |
| | max_features | {sqrt, log2, None} |
| | splitter | {best, random} |
| | criterion | {squared_error, friedman_mse, absolute_error, poisson} |
| **RandomForest / ExtraTrees** | n_estimators | $50, 500$ |
| | max_depth | $3, 30$ |
| | min_samples_split | $2, 20$ |
| | min_samples_leaf | $1, 10$ |
| | max_features | {sqrt, log2, None} |
| | bootstrap | {True, False} |
| **AdaBoost** | n_estimators | $50, 500$ |
| | learning_rate | $0.001, 2.0$ (log) |
| | loss | {linear, square, exponential} |
| **GradientBoosting** | n_estimators | $50, 500$ |
| | learning_rate | $0.001, 0.3$ (log) |
| | max_depth | $2, 10$ |
| | min_samples_split | $2, 20$ |
| | min_samples_leaf | $1, 10$ |
| | subsample | $0.5, 1.0$ |
| | max_features | {sqrt, log2, None} |
| | loss | {squared_error, absolute_error, huber} |
| **HistGradientBoosting** | max_iter | $50, 500$ |
| | learning_rate | $0.001, 0.3$ (log) |
| | max_depth | $2, 15$ |
| | min_samples_leaf | $5, 50$ |
| | l2_regularization | $0.0, 10.0$ |
| | max_bins | $128, 255$ |
| | loss | {squared_error, absolute_error} |
| *Gradient Boosting Libraries* | | |
| **CatBoost** | iterations | $50, 500$ |
| | learning_rate | $0.001, 0.3$ (log) |
| | depth | $2, 10$ |
| | l2_leaf_reg | $1.0, 10.0$ (log) |
| | border_count | $32, 255$ |
| | random_strength | $0.0, 10.0$ |
| | bagging_temperature | $0.0, 1.0$ |
| **XGBoost** | n_estimators | $50, 500$ |
| | learning_rate | $0.001, 0.3$ (log) |
| | max_depth | $2, 10$ |
| | min_child_weight | $1, 10$ |
| | subsample | $0.5, 1.0$ |
| | colsample_bytree | $0.5, 1.0$ |
| | gamma | $0.0, 5.0$ |
| | reg_alpha | $0.0, 1.0$ |
| | reg_lambda | $0.0, 1.0$ |
| *Bagging Wrappers* | | |
| **BaggingSVR** | C | $0.1, 100$ (log) |
| | epsilon | $0.01, 1.0$ (log) |
| | kernel | {rbf} |
| | gamma | {scale, auto} |
| **BaggingDecisionTree** | max_depth | $3, 20$ |

*Continued on next page*

| Pipeline | Hyperparameter | Search Space |
|---|---|---|
| | min_samples_split | $2, 20$ |
| | min_samples_leaf | $1, 10$ |
| | max_features | {sqrt, log2, None} |
| **BaggingMLP** | hidden_layer_sizes | {(64,), (128,), (64,32), (128,64), (128,64,32)} |
| | activation | {relu, tanh} |
| | alpha | $10^{-5}, 10^{-1}$ (log) |
| | learning_rate_init | $10^{-4}, 10^{-2}$ (log) |
| *Deep Learning* | | |
| **Deep Learning** | learning_rate | $\{$1e-5, 5e-5, 1e-4, 5e-4, 1e-3, 5e-3$\}$ |
| | weight_decay | $\{$0, 1e-5, 1e-4, 1e-3, 1e-2$\}$ |
| | dropout | $\{$0.0, 0.1, 0.2, 0.3, 0.4, 0.5$\}$ |
| | batch_size | $\{$32, 64, 128, 256$\}$ |
| | optimizer | {adam, adamw, radam} |
| | loss_type | {mse, huber, gaussian_nll, log_cosh} |
| | activation | {ReLU, GELU, ELU, LeakyReLU, SiLU} |
| | scheduler | {warmup_cosine, cosine, plateau, onecycle} |
| | warmup_epochs | $\{0, 3, 5, 10\}$ |
| | gradient_clip | {None, 0.5, 1.0, 5.0} |
| | early_stopping_patience | $\{10, 15, 20, 30\}$ |
| | scaler_type | {standard, robust, none} |

*Note:* Architecture-specific hyperparameters (e.g., number of layers, hidden dimensions, attention heads) are tuned per model and described in the supplementary materials. BaggingSVR, BaggingDecisionTree, and BaggingMLP Extreme variants share identical base-estimator search spaces; they differ only in the sigmoid bootstrap sampling dynamic range (100$\times$ vs. 10$\times$).

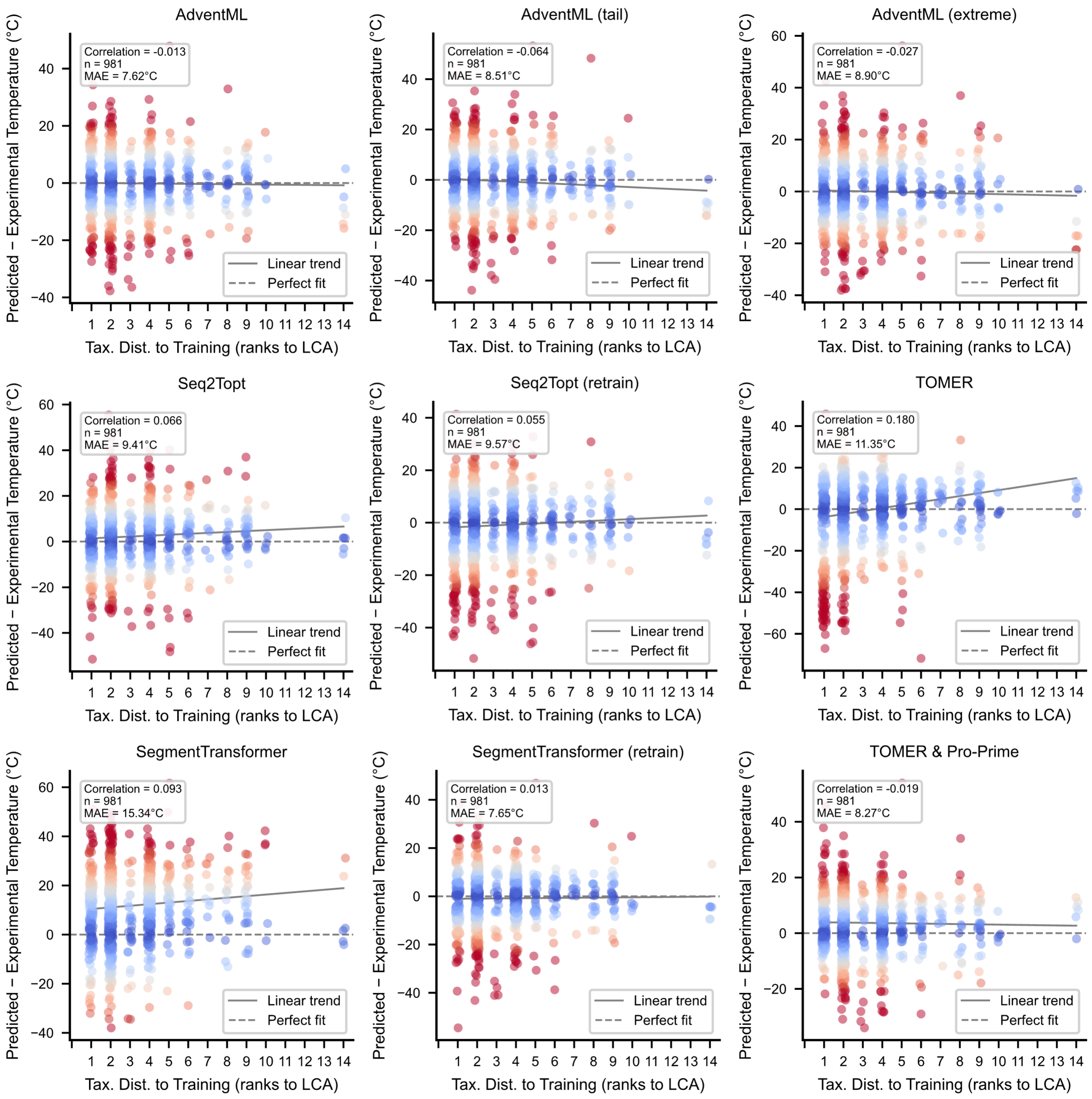


**Figure S2. Prediction error stratified by phylogenetic distance.** Model performance as a function of phylogenetic distance between test and training sequences. Phylogenetic distance showed by branch count between test observation, closest training sample and last common ancestor. Positive values correspond to predictions exceeding the experimental $T_{opt}$.

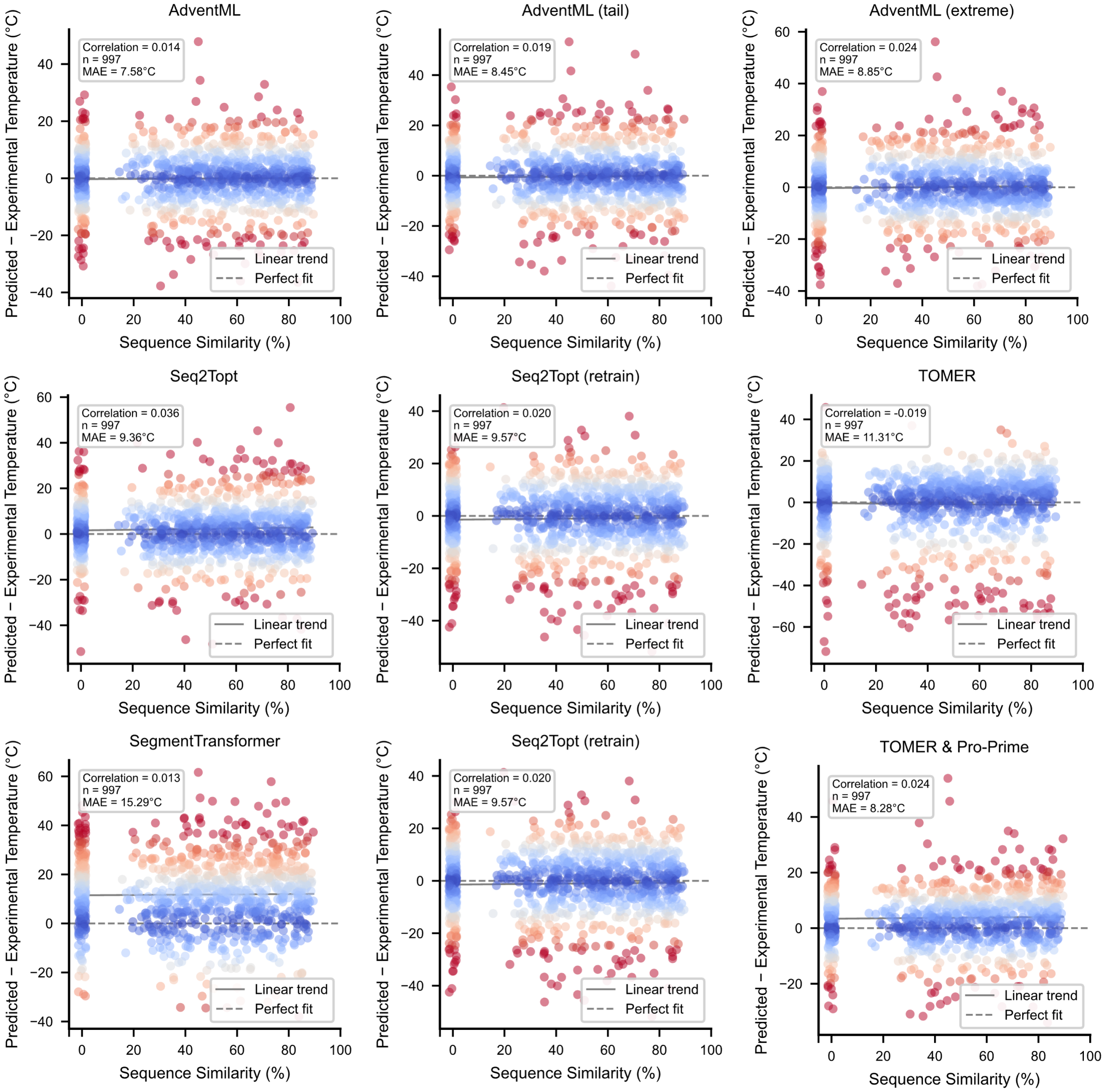


**Figure S3. Prediction performance as a function of sequence similarity.** Model accuracy stratified by maximum sequence identity between each test sequence and the training set. Positive values correspond to predictions exceeding the experimental $T_{opt}$.

**Table S10.** Protein representations names and descriptions.

| Abbreviation | Full Written Name / Description | Category |
|---|---|---|
| PT T5 XL U50 | ProtTrans T5-XL-U50 | Transformer |
| PT T5 U50 | ProtTrans T5-UniRef50 | Transformer |
| PT T5 BFD | ProtTrans T5-BFD | Transformer |
| PT Bert BFD | ProtTrans BERT-BFD | Transformer |
| PT Albert BFD | ProtTrans ALBERT-BFD | Transformer |
| PT XLNet U100 | ProtTrans XLNet-UniRef100 | Transformer |
| ProstT5 | RostLab ProstT5 | Transformer |
| ESM2 35B | ESM-2 (35 Billion parameters) | Transformer |
| ESM2 3B | ESM-2 (3 Billion parameters) | Transformer |
| ESM2 650M | ESM-2 (650 Million parameters) | Transformer |
| ESM2 35M | ESM-2 (35 Million parameters) | Transformer |
| ESM2 | ESM-2 (Base/Default version) | Transformer |
| ESM1b | ESM-1b | Transformer |
| ESMC 600M | ESM-C (600 Million parameters) | Transformer |
| ESMC 300M | ESM-C (300 Million parameters) | Transformer |
| ESM3 SM | ESM-3 Small Open v1 | Transformer |
| Ankh3 XL | Ankh3 Extra Large | Transformer |
| Ankh3 Large | Ankh3 Large | Transformer |
| Ankh Large | Ankh Large | Transformer |
| Ankh Base | Ankh Base | Transformer |
| Prime | AI4Protein/Prime | Transformer |
| PLUS RNN | PLUS Recurrent Neural Network | Other Learned |
| SeqVec | Sequence Vector (SeqVec) | Other Learned |
| GloVe | Global Vectors for Word Representation | Other Learned |
| FastText | FastText | Other Learned |
| CPCProt | Contrastive Predictive Coding for Proteins | Other Learned |
| Bepler | Bepler & Berger Embeddings | Other Learned |
| *Physicochemical | **Comprehensive Physicochemical Properties:** Includes molecular weight, hydrophobicity, isoelectric point, normalized van der Waals volume, polarizability, generic structural descriptors, and solvent accessibility metrics. | Physicochemical |
| *Composition | **Amino Acid & PAAC:** Amino acid frequency composition (20 features) and Pseudo-amino acid composition (PAAC), which incorporates sequence order information. | Physicochemical |
| *AAIndex | **AAindex-derived Properties:** Direct numerical indices of all physicochemical and biological properties of amino acids (AAindex database). | Physicochemical |
| *Auto | **Autocorrelation Descriptors:** Moreau-Broto, Moran, and Geary autocorrelation metrics that measure the distribution of amino acid properties along the protein sequence. | Physicochemical |
| *Simple | **Simple Descriptors & CTD:** Conjoint Triad Descriptors (CTD) counting the continuous sequences of three amino acids grouped by dipole scale and volume, alongside basic sequence length bounds. | Physicochemical |

### Pseudocode for Resampling Strategies

The following pseudocode describes the resampling algorithms implemented in this work for imbalanced regression tasks. All algorithms operate on a dataset $DX, y$ with features $X$ and continuous target values $y$.

#### Oversampling Methods

**Algorithm 1** ADASYN Regressor (Adaptive Synthetic Sampling)

**Require:** $DX, y$ – dataset with features $X$ and continuous target $y$  
**Require:** $k$ – number of nearest neighbors  
**Require:** $t_R$ – relevance threshold (quantile on $|y -$ median$y|$)  
**Require:** $c$ – relevance coefficient (multiplier for synthetic samples)  
**Require:** *method* – sampling method: 'balance' or 'extreme'  
**Ensure:** *newD* – resampled dataset  
$d_i \leftarrow |y_i -$ median$y|$ for all instances  
$\tau \leftarrow$ quantile$d, t_R$  
*extremeD* $\leftarrow$ instances in $D$ where $d_i \geq \tau$  
*nonExtremeD* $\leftarrow$ instances in $D$ where $d_i < \tau$  
**if** *method* = 'balance' **then**  
    $G \leftarrow |$*nonExtremeD*$| - |$*extremeD*$| \times c$  
**else**  
    $G \leftarrow |$*extremeD*$| \times c$  
**end if**  
*nns* $\leftarrow$ get $k$-nearest neighbors for all instances in *extremeD*  
**for** each instance $x_i$ in *extremeD* **do**  
    $r_i \leftarrow$ fraction of neighbors that are non-extreme  
**end for**  
$\hat{r}_i \leftarrow r_i r_i$ ▷ normalize  
$g_i \leftarrow$ round$\hat{r}_i \times G$ ▷ allocate synthetic samples per instance  
*syntheticD* $\leftarrow \emptyset$  
**for** each instance $x_i$ in *extremeD* with allocation $g_i$ **do**  
    **for** $j \leftarrow 1$ to $g_i$ **do**  
        $x_{nb} \leftarrow$ random neighbor from *nnsi*  
        $\lambda \leftarrow$ random$0, 1$  
        $x_{syn} \leftarrow x_i \; \lambda \times x_{nb} - x_i$  
        $y_{syn} \leftarrow y_i \; \lambda \times y_{nb} - y_i$  
        *syntheticD* $\leftarrow$ *syntheticD* $\cup \{x_{syn}, y_{syn}\}$  
    **end for**  
**end for**  
**return** $D \cup$ *syntheticD*

#### Undersampling Methods

#### Extreme-Only Methods

The following methods are implemented only with fixed tail thresholds, as they are simple sampling strategies that benefit from domain-specific region definitions.

#### Extreme Variants of Adaptive Methods

The extreme variants of all above adaptive algorithms use **fixed tail thresholds** instead of adaptive/histogram-based detection of rare regions. Specifically:

- **Tail region**: instances where $y < 25$ or $y > 60$
- **Middle region**: instances where $25 \leq y \leq 60$

**Key differences between standard and extreme variants:**

1. **Standard variants**: Use adaptive thresholds based on:
    - Histogram binning (SMOTER, SMOGN, KNNOR, NearMiss)

**Algorithm 2** SMOTER Regressor (SMOTE for Regression)

**Require:** $DX, y$ – dataset with features $X$ and continuous target $y$
**Require:** $s$ – sampling strategy (number or fraction of synthetic samples)
**Require:** $k$ – number of nearest neighbors
**Require:** $b$ – number of bins (default: $\sqrt{n}$)
**Ensure:** *newD* – resampled dataset
*hist*, *edges* $\leftarrow$ histogram$y, b$
*avg* $\leftarrow$ mean*hist*
*rareBins* $\leftarrow$ bins where *hist*$i <$ *avg*
*rareD* $\leftarrow$ instances in $D$ falling in *rareBins*
$m \leftarrow |$*rareD*$|$
**if** $s$ is float **then**
    $n_{synth} \leftarrow \lfloor s \times m \rfloor$
**else**
    $n_{synth} \leftarrow s$
**end if**
*nns* $\leftarrow$ get $k$-nearest neighbors for all instances in *rareD*
*syntheticD* $\leftarrow \emptyset$
**while** $|$*syntheticD*$| < n_{synth}$ **do**
    $i \leftarrow$ random index in *rareD*
    $j \leftarrow$ random neighbor from *nns*$i$
    $u \leftarrow$ random$0, 1$
    $x_{syn} \leftarrow x_i \; u \times x_j - x_i$
    $y_{syn} \leftarrow y_i \; u \times y_j - y_i$
    *syntheticD* $\leftarrow$ *syntheticD* $\cup \{x_{syn}, y_{syn}\}$
**end while**
**return** $D \cup$ *syntheticD*

**Algorithm 3** SMOGN Regressor (SMOTE with Gaussian Noise)

**Require:** $DX, y$ – dataset with features $X$ and continuous target $y$
**Require:** $s$ – sampling strategy (number or fraction of synthetic samples)
**Require:** $k$ – number of nearest neighbors
**Require:** $b$ – number of bins (default: $\sqrt{n}$)
**Require:** $\sigma$ – standard deviation for Gaussian noise
**Ensure:** *newD* – resampled dataset
*hist*, *edges* $\leftarrow$ histogram$y, b$
*avg* $\leftarrow$ mean*hist*
*rareBins* $\leftarrow$ bins where *hist*$i <$ *avg*
*rareD* $\leftarrow$ instances in $D$ falling in *rareBins*
$m \leftarrow |$*rareD*$|$
**if** $s$ is float **then**
    $n_{synth} \leftarrow \lfloor s \times m \rfloor$
**else**
    $n_{synth} \leftarrow s$
**end if**
*nns* $\leftarrow$ get $k$-nearest neighbors for all instances in *rareD*
*syntheticD* $\leftarrow \emptyset$
**while** $|$*syntheticD*$| < n_{synth}$ **do**
    $i \leftarrow$ random index in *rareD*
    $j \leftarrow$ random neighbor from *nns*$i$
    $u \leftarrow$ random$0, 1$
    $x_{syn} \leftarrow x_i \; u \times x_j - x_i \; \mathcal{N}0, \sigma$
    $y_{syn} \leftarrow y_i \; u \times y_j - y_i \; \mathcal{N}0, \sigma$
    *syntheticD* $\leftarrow$ *syntheticD* $\cup \{x_{syn}, y_{syn}\}$
**end while**
**return** $D \cup$ *syntheticD*

**Algorithm 4** WERCS Regressor (Weighted Relevance Combination Sampling)

**Require:** $DX, y$ – dataset with features $X$ and continuous target $y$
**Require:** $s$ – sampling strategy (number or fraction of synthetic samples)
**Require:** $k$ – number of nearest neighbors
**Ensure:** *newD* – resampled dataset
$n \leftarrow |D|$
**if** $s$ is float **then**
    $n_{synth} \leftarrow \lfloor s \times n \rfloor$
**else**
    $n_{synth} \leftarrow s$
**end if**
*med* $\leftarrow$ median$y$
$w_i \leftarrow |y_i - \mathit{med}|$ for all instances ▷ weight by deviation from median
$\hat{w}_i \leftarrow w_i\, w_i$ ▷ normalize weights
$c_i \leftarrow \lfloor \hat{w}_i \times n_{synth} \rfloor$ ▷ allocate counts per instance
Adjust $c_i$ to ensure $c_i = n_{synth}$
*nns* $\leftarrow$ get $k$-nearest neighbors for all instances in $D$
*syntheticD* $\leftarrow \emptyset$
**for** each instance $x_i$ in $D$ with allocation $c_i$ **do**
    **for** $j \leftarrow 1$ to $c_i$ **do**
        $x_{nb} \leftarrow$ random neighbor from *nns*$i$
        $u \leftarrow$ random$0, 1$
        $x_{syn} \leftarrow x_i\; u \times x_{nb} - x_i$
        $y_{syn} \leftarrow y_i\; u \times y_{nb} - y_i$
        *syntheticD* $\leftarrow$ *syntheticD* $\cup \{x_{syn}, y_{syn}\}$
    **end for**
**end for**
**return** $D \cup$ *syntheticD*

**Algorithm 5** WERCS-GN Regressor (WERCS with Gaussian Noise)

**Require:** $DX, y$ – dataset with features $X$ and continuous target $y$
**Require:** $s$ – sampling strategy (number or fraction of synthetic samples)
**Require:** $k$ – number of nearest neighbors
**Require:** $\sigma$ – standard deviation for Gaussian noise
**Ensure:** *newD* – resampled dataset

$n \leftarrow |D|$
**if** $s$ is float **then**
  $n_{synth} \leftarrow \lfloor s \times n \rfloor$
**else**
  $n_{synth} \leftarrow s$
**end if**
*med* $\leftarrow$ median$y$
$w_i \leftarrow |y_i - med|$ for all instances
$\hat{w}_i \leftarrow w_i w_i$
$c_i \leftarrow \lfloor \hat{w}_i \times n_{synth} \rfloor$
Adjust $c_i$ to ensure $c_i = n_{synth}$
*nns* $\leftarrow$ get $k$-nearest neighbors for all instances in $D$
*syntheticD* $\leftarrow \emptyset$
**for** each instance $x_i$ in $D$ with allocation $c_i$ **do**
  **for** $j \leftarrow 1$ to $c_i$ **do**
    $x_{nb} \leftarrow$ random neighbor from *nns*$i$
    $u \leftarrow$ random$0, 1$
    $x_{syn} \leftarrow x_i \; u \times x_{nb} - x_i \; \mathcal{N}0, \sigma$
    $y_{syn} \leftarrow y_i \; u \times y_{nb} - y_i \; \mathcal{N}0, \sigma$
    *syntheticD* $\leftarrow$ *syntheticD* $\cup \{x_{syn}, y_{syn}\}$
  **end for**
**end for**
**return** $D \cup$ *syntheticD*

**Algorithm 6** KNNOR Regressor (K-Nearest Neighbors Oversampling for Regression)

**Require:** $DX, y$ – dataset with features $X$ and continuous target $y$
**Require:** $k$ – number of nearest neighbors
**Require:** $p$ – proportion of minority for threshold determination
**Require:** $\alpha$ – maximum interpolation factor
**Require:** $b$ – number of bins (default: $\sqrt{n}$)
**Require:** $t$ – target count per bin (default: mean bin frequency)
**Ensure:** *newD* – resampled dataset

*hist*, *edges* $\leftarrow$ histogram$y, b$
**if** $t$ = None **then**
    $t \leftarrow$ mean*hist*
**end if**
*syntheticD* $\leftarrow \emptyset$
**for** each bin $i$ where *hist*$i < t$ **do**
    $D_{bin} \leftarrow$ instances in $D$ falling in bin $i$
    *toAdd* $\leftarrow t - |D_{bin}|$
    Compute $k$-th neighbor distances for all instances in $D_{bin}$
    $\tau \leftarrow$ threshold at $p$-th percentile of distances
    **while** *toAdd* $> 0$ **do**
        **for** each $x_i$ in $D_{bin}$ where $k$-th neighbor distance $\leq \tau$ **do**
            $x_{syn} \leftarrow x_i$
            **for** $j \leftarrow 1$ to $k$ **do**
                $x_{nb} \leftarrow j$-th neighbor of $x_i$
                $\lambda \leftarrow$ random$0, \alpha$
                $x_{syn} \leftarrow x_{syn}\ \lambda \times x_{nb} - x_i$
            **end for**
            $y_{syn} \leftarrow$ inverse-distance weighted average of neighbor $y$ values
            *syntheticD* $\leftarrow$ *syntheticD* $\cup \{x_{syn}, y_{syn}\}$
            *toAdd* $\leftarrow$ *toAdd* $- 1$
            **if** *toAdd* $= 0$ **then**
                **break**
            **end if**
        **end for**
    **end while**
**end for**
**return** $D \cup$ *syntheticD*

**Algorithm 7** Cluster Centroids Regressor

**Require:** $DX, y$ – dataset with features $X$ and continuous target $y$
**Require:** $s$ – sampling strategy (desired number or fraction of samples)
**Require:** *voting* – strategy: 'hard' or 'soft'
**Ensure:** *newD* – resampled dataset
$n \leftarrow |D|$
**if** $s$ is float **then**
  $n_{samples} \leftarrow \lfloor s \times n \rfloor$
**else**
  $n_{samples} \leftarrow s$
**end if**
Fit KMeans clustering with $n_{clusters} = n_{samples}$
*centers* $\leftarrow$ cluster centroids
**if** *voting* = 'hard' **then**
  Find nearest original sample to each centroid
  $X_{res} \leftarrow$ nearest samples' features
  $y_{res} \leftarrow$ nearest samples' targets
**else** ▷ soft voting
  $X_{res} \leftarrow$ *centers*
  $y_{res}i \leftarrow$ mean$y_j$ for all $j$ assigned to cluster $i$
**end if**
**return** $X_{res}, y_{res}$

**Algorithm 8** CNN Regressor (Condensed Nearest Neighbour)

**Require:** $DX, y$ – dataset with features $X$ and continuous target $y$
**Require:** $s$ – sampling strategy (desired number or fraction of samples)
**Require:** $k$ – number of neighbors for KNN regressor
**Require:** $n_{seeds}$ – number of initial seed samples
**Require:** $\epsilon$ – tolerance threshold for prediction error
**Ensure:** *newD* – resampled dataset
$n \leftarrow |D|$
**if** $s$ is float **then**
  $n_{desired} \leftarrow \lfloor s \times n \rfloor$
**else**
  $n_{desired} \leftarrow s$
**end if**
$C \leftarrow$ randomly select $n_{seeds}$ instances from $D$ ▷ condensed set
*changed* $\leftarrow$ True
**while** *changed* **do**
  *changed* $\leftarrow$ False
  Train KNN regressor on $C$
  **for** each instance $x_i, y_i$ in $D \setminus C$ **do**
    $\hat{y}_i \leftarrow$ KNN prediction for $x_i$
    **if** $|y_i - \hat{y}_i| > \epsilon$ **then**
      $C \leftarrow C \cup \{x_i, y_i\}$
      *changed* $\leftarrow$ True
    **end if**
  **end for**
**end while**
**if** $|C| > n_{desired}$ **then**
  $C \leftarrow$ random subset of $C$ with $n_{desired}$ instances
**end if**
**return** $C$

**Algorithm 9** ENN Regressor (Edited Nearest Neighbours)

**Require:** $DX, y$ – dataset with features $X$ and continuous target $y$
**Require:** $s$ – sampling strategy (desired number or fraction of samples)
**Require:** $k$ – number of neighbors
**Require:** $\epsilon$ – tolerance threshold for prediction error
**Ensure:** *newD* – resampled dataset

$n \leftarrow |D|$
**if** $s$ is float **then**
    $n_{\textit{desired}} \leftarrow \lfloor s \times n \rfloor$
**else**
    $n_{\textit{desired}} \leftarrow s$
**end if**
Train KNN regressor on $D$
**for** each instance $x_i, y_i$ in $D$ **do**
    *neighbors* $\leftarrow k$ nearest neighbors of $x_i$ (excluding self)
    $\hat{y}_i \leftarrow$ mean$y_j$ for $j$ in *neighbors*
    $e_i \leftarrow |y_i - \hat{y}_i|$
**end for**
*keepIndices* $\leftarrow$ indices where $e_i \leq \epsilon$
**if** $|$*keepIndices*$| > n_{\textit{desired}}$ **then**
    *keepIndices* $\leftarrow$ random subset with $n_{\textit{desired}}$ indices
**end if**
**return** instances in $D$ at *keepIndices*

**Algorithm 10** IHT Regressor (Instance Hardness Threshold)

**Require:** $DX, y$ – dataset with features $X$ and continuous target $y$
**Require:** $s$ – sampling strategy (desired number or fraction of samples)
**Require:** *cv* – number of cross-validation folds
**Require:** *estimator* – regression model (default: RandomForest)
**Ensure:** *newD* – resampled dataset

$n \leftarrow |D|$
**if** $s$ is float **then**
    $n_{\textit{desired}} \leftarrow \lfloor s \times n \rfloor$
**else**
    $n_{\textit{desired}} \leftarrow s$
**end if**
$\hat{y} \leftarrow$ cross-validated predictions using *estimator* with *cv* folds
$e_i \leftarrow |y_i - \hat{y}_i|$ for all instances ▷ instance hardness
*sortedIndices* $\leftarrow$ argsort$e$ ▷ sort by ascending error
*selected* $\leftarrow$ *sortedIndices*$1 : n_{\textit{desired}}$ ▷ keep easiest instances
**return** instances in $D$ at *selected*

**Algorithm 11** NearMiss Regressor

**Require:** $DX, y$ – dataset with features $X$ and continuous target $y$
**Require:** $s$ – sampling strategy (number or fraction of dense-region samples to retain)
**Require:** *version* – NearMiss version: 1, 2, or 3
**Require:** $k$ – number of neighbors
**Require:** $b$ – number of bins (default: $\sqrt{n}$)
**Ensure:** *newD* – resampled dataset

*hist*, *edges* $\leftarrow$ histogram$y, b$
*avg* $\leftarrow$ mean*hist*
*sparseBins* $\leftarrow$ bins where *hist*$i \leq$ *avg*
*denseBins* $\leftarrow$ bins where *hist*$i >$ *avg*
*sparseD* $\leftarrow$ instances falling in *sparseBins*
*denseD* $\leftarrow$ instances falling in *denseBins*
**if** $s$ is float **then**
    $n_{keep} \leftarrow \lfloor s \times |\mathit{denseD}| \rfloor$
**else**
    $n_{keep} \leftarrow s$
**end if**
**if** *version* $= 1$ **then**
    For each $x_i$ in *denseD*: compute mean distance to $k$ nearest sparse neighbors
    *selected* $\leftarrow n_{keep}$ instances with smallest mean distance
**else if** *version* $= 2$ **then**
    For each $x_i$ in *denseD*: compute mean distance to all sparse instances
    *selected* $\leftarrow n_{keep}$ instances with smallest mean distance
**else** ▷ version 3
    For each $x_i$ in *sparseD*: find $k$ nearest dense neighbors
    *candidates* $\leftarrow$ unique dense neighbors found
    For each $x_i$ in *candidates*: compute mean distance to $k$ nearest sparse neighbors
    *selected* $\leftarrow n_{keep}$ candidates with largest mean distance
**end if**
**return** instances in *denseD* at *selected*

**Algorithm 12** OSS Regressor (One-Sided Selection)

**Require:** $DX, y$ – dataset with features $X$ and continuous target $y$
**Require:** $s$ – sampling strategy (number or fraction of non-critical samples to retain)
**Require:** $p$ – critical percentile (defines lower and upper tails)
**Require:** $k$ – number of neighbors for KNN
**Require:** $n_{seeds}$ – number of initial seeds from non-critical region
**Ensure:** *newD* – resampled dataset

*low* $\leftarrow$ percentile$y, p \times 100$
*high* $\leftarrow$ percentile$y, 100 - p \times 100$
*criticalD* $\leftarrow$ instances where $y \leq$ *low* or $y \geq$ *high*
*nonCritD* $\leftarrow$ instances where *low* $< y <$ *high*
**if** $s$ is float **then**
    $n_{keep} \leftarrow \lfloor s \times |\mathit{nonCritD}| \rfloor$
**else**
    $n_{keep} \leftarrow s$
**end if**
*seeds* $\leftarrow$ random selection of $n_{seeds}$ instances from *nonCritD*
$C \leftarrow$ *criticalD* $\cup$ *seeds* ▷ initial condensed set
Train KNN regressor on $C$
*remaining* $\leftarrow$ *nonCritD* $\setminus$ *seeds*
**for** each $x_i, y_i$ in *remaining* **do**
    $e_i \leftarrow |y_i -$ KNN.predict$x_i|$
**end for**
*selectedNonCrit* $\leftarrow n_{keep} - n_{seeds}$ instances with largest $e_i$
**return** *criticalD* $\cup$ *seeds* $\cup$ *selectedNonCrit*

**Algorithm 13** Tomek Links Regressor

**Require:** $DX, y$ – dataset with features $X$ and continuous target $y$
**Require:** $p$ – threshold percentile for target difference
**Ensure:** *newD* – resampled dataset
*nn* $\leftarrow$ 1-nearest neighbor for all instances in $D$
*mutualPairs* $\leftarrow \emptyset$
**for** each instance $i$ in $D$ **do**
  $j \leftarrow$ nearest neighbor of $i$
  **if** nearest neighbor of $j$ is $i$ **then** ▷ mutual nearest neighbors
    *mutualPairs* $\leftarrow$ *mutualPairs* $\cup \{i, j\}$ where $i < j$
  **end if**
**end for**
*diffs* $\leftarrow |y_i - y_j|$ for all $i, j$ in *mutualPairs*
$\tau \leftarrow$ percentile*diffs*$, p$
*toRemove* $\leftarrow \emptyset$
**for** each $i, j, d$ in *mutualPairs*, *diffs* **do**
  **if** $d \geq \tau$ **then** ▷ Tomek link with large target discrepancy
    *toRemove* $\leftarrow$ *toRemove* $\cup \{i, j\}$
  **end if**
**end for**
**return** $D \setminus$ *toRemove*

**Algorithm 14** Extreme Random Oversampling Regressor

**Require:** $DX, y$ – dataset with features $X$ and continuous target $y$
**Require:** $s$ – sampling strategy (number or fraction of samples to duplicate)
**Require:** $\tau_{low} = 25$ – lower tail threshold
**Require:** $\tau_{high} = 60$ – upper tail threshold
**Ensure:** *newD* – resampled dataset
*tailD* $\leftarrow$ instances where $y < \tau_{low}$ or $y > \tau_{high}$
$m \leftarrow |$*tailD*$|$
**if** $m = 0$ **then**
  **return** $D$
**end if**
**if** $s$ is float **then**
  $n_{synth} \leftarrow \lfloor s \times m \rfloor$
**else**
  $n_{synth} \leftarrow s$
**end if**
*duplicatedIdx* $\leftarrow$ randomly sample $n_{synth}$ indices from *tailD* with replacement
$X_{syn} \leftarrow X$*duplicatedIdx*
$y_{syn} \leftarrow y$*duplicatedIdx*
**return** $D \cup X_{syn}, y_{syn}$

**Algorithm 15** Extreme Random Undersampling Regressor

**Require:** $DX, y$ – dataset with features $X$ and continuous target $y$
**Require:** $s$ – sampling strategy (number or fraction of middle samples to retain)
**Require:** $\tau_{low} = 25$ – lower tail threshold
**Require:** $\tau_{high} = 60$ – upper tail threshold
**Ensure:** *newD* – resampled dataset
*tailD* $\leftarrow$ instances where $y < \tau_{low}$ or $y > \tau_{high}$
*middleD* $\leftarrow$ instances where $\tau_{low} \leq y \leq \tau_{high}$
$m \leftarrow |\textit{middleD}|$
**if** $m = 0$ **then**
    **return** $D$
**end if**
**if** $s$ is float **then**
    $n_{keep} \leftarrow \lfloor s \times m \rfloor$
**else**
    $n_{keep} \leftarrow s$
**end if**
*keepIdx* $\leftarrow$ randomly sample $n_{keep}$ indices from *middleD* without replacement
*middleD*$_{res}$ $\leftarrow$ instances at *keepIdx*
**return** *tailD* $\cup$ *middleD*$_{res}$ ▷ all tail samples preserved

**Algorithm 16** Extreme Variant Modification

**Require:** $DX, y$ – dataset with features $X$ and continuous target $y$
**Require:** $\tau_{low} = 25$ – lower tail threshold
**Require:** $\tau_{high} = 60$ – upper tail threshold
**Ensure:** Region identification for extreme variants
*tailD* $\leftarrow$ instances where $y < \tau_{low}$ or $y > \tau_{high}$
*middleD* $\leftarrow$ instances where $\tau_{low} \leq y \leq \tau_{high}$
**For oversampling methods** (ADASYN, SMOTER, SMOGN, WERCS, WERCS-GN, KNNOR):
    Generate synthetic samples **only from** *tailD*
    All original samples in $D$ are preserved
**For undersampling methods** (ClusterCentroids, CNN, ENN, IHT, NearMiss, OSS, TomekLinks):
    Apply undersampling **only to** *middleD*
    All samples in *tailD* are preserved unchanged

- Quantile-based relevance thresholds (ADASYN)
- Percentile-based critical regions (OSS)
- Prediction error thresholds (CNN, ENN, IHT)
- Target discrepancy percentiles (TomekLinks)
- Deviation from median (WERCS, WERCS-GN)

2. **Extreme variants**: Use fixed domain-specific thresholds ($y < 25$ or $y > 60$) to identify samples requiring special handling, ensuring that tail regions are always prioritized regardless of the dataset's distribution.
